# Metabolic tuning facilitates nociceptor resilience to excitotoxicity

**DOI:** 10.1101/2024.12.23.630176

**Authors:** Lin Yuan, Navdeep S. Chandel, David Julius

## Abstract

The capsaicin receptor, TRPV1, mediates the detection of harmful chemical and thermal stimuli. Overactivation of TRPV1 can lead to cellular damage or death through excitotoxicity, a phenomenon associated with painful neuropathy and the paradoxical use of capsaicin as an analgesic. We exploited capsaicin-evoked death to conduct a systematic analysis of excitotoxicity through a genome-wide CRISPRi screen, thereby revealing a comprehensive network of regulatory pathways. We show that decreased expression of mitochondrial electron transport chain (ETC) components protects against capsaicin-induced toxicity by mitigating calcium imbalance and mitochondrial reactive oxygen species production via distinct pathways. Interestingly, TRPV1^+^ sensory neurons in adult mice maintain lower expression of ETC components and can better tolerate excitotoxicity and oxidative stress compared to other sensory neuron subtypes. We further confirm the regulatory roles of the ETC in sensory neurons through gain-of-function and loss-of-function experiments. These findings implicate ETC tuning as a cellular protective strategy against sensory excitotoxicity.

## INTRODUCTION

Excitotoxicity describes the phenomenon of neuronal damage or death caused by overactivation of excitatory receptors leading to calcium overload. It is traditionally and most extensively studied in the context of glutamate toxicity in the central nervous system (CNS), which is thought to mediate cell death following ischemia and cell injury in many neurodegenerative diseases including Alzheimer’s and Parkinson’s disease.^1,2^ Although excitotoxicity is associated with many disease conditions in the peripheral nervous system (PNS), how it is mediated in PNS neurons and how it contributes to peripheral neural pathology are largely unexplored.

Primary afferent nociceptors are a subset of peripheral sensory neurons that specialize in detecting noxious stimuli and propagating the signal to initiate pain sensation. Intense, repetitive, or prolonged excitation of these neurons can result in chronic and maladaptive pain as well as afferent degeneration. For example, studies of hereditary painful neuropathies revealed gain-of-function mutations in excitatory ion channels, including voltage-gated sodium (NaV1.7, NaV1.8, NaV1.9) or TRPV4 channels, that can render nociceptors hyperexcitable and degenerative.^3-6^ Similarly, hyper-excitability and degeneration are also observed in more common forms of painful neuropathy, such as diabetic neuropathy,^7,8^ but whether hyper-excitability leads to excitotoxicity and degeneration is less examined. This is likely due to the lack of mechanistic understanding of excitotoxicity in the peripheral nociceptive neurons, which is needed to better characterize these prevalent and insidious diseases and aid in the development of therapeutic options, which are currently lacking.

Capsaicin, the pungent ingredient of chili peppers, has long been recognized as an excitatory neurotoxin for sensory neurons.^9,10^ Following prolonged or repetitive exposure, capsaicin selectively injures or kills cells that express its receptor, the excitatory TRPV1 ion channel,^11^ which is a defining molecular marker for a major class of nociceptors. In fact, the paradoxical use of capsaicin creams or patches as non-opioid analgesics is based on the finding that topical capsaicin application selectively decreases TRPV1+ epidermal nerve fiber density, which can provide pain relief until these terminals regenerate.^12^ This highlights the major roles TRPV1+ afferents play in initiating painful sensation as well as the specific and robust neurotoxic action of capsaicin.

As an non-selective cation channel, TRPV1 is activated by noxious chemical and thermal stimuli and its activation can be potentiated by inflammatory mediators, such as extracellular protons, peptides, and bioactive lipids, that drive pain hypersensitivity and nerve injury.^13,14^ TRPV1 is also highly permeable to calcium ions,^11,15^ and, in this way, analogous to NMDA glutamate receptors, which play a key role in excitotoxicity of CNS neurons.^1,2^ Recent evidence suggests that TRPV1-mediated excitotoxicity underlies dry-eye-disease associated corneal nerve damage,^16^ underscoring the need to further delineate mechanisms underlying this form of peripheral excitotoxicity. Interestingly, human or even yeast cells can acquire vulnerability to capsaicin when they heterologously express TRPV1,^11,17^ suggesting that the mechanism of excitotoxicity involves a conserved cell death pathway. Moreover, this observation demonstrates that such heterologous expression systems can be leveraged as facile cellular models to probe mechanisms of excitotoxicity.

With this in mind, we designed an unbiased, genome-wide CRISPRi screen to identify molecules and pathways that enhance or diminish capsaicin-evoked death in immortalized mammalian cells. Foremost among these ‘hits’ are components of the mitochondrial electron transport chain (ETC), suppression of which protects against capsaicin-induced excitotoxicity in both TRPV1-expressing (TRPV1+) cell lines and adult mouse sensory neurons. Pharmacologic and genetic experiments suggest that ETC suppression boosts cell resilience by two parallel pathways: reducing mitochondrial reactive oxygen species (ROS) and hence oxidative stress and regulating TRPV1 localization. Furthermore, we find that TRPV1+ nociceptors from adult mice express ETC components at lower levels compared to other sensory neurons, conferring greater resilience to cell death due to calcium or ROS overload. We therefore propose that tuning of aerobic respiration helps nociceptors mitigate risk associated with injury and other noxious events. These insights into cellular mechanisms controlling nociceptor excitotoxicity are relevant to understanding neuropathological consequences of diabetes, chemotherapy, and other conditions that disrupt normal pain sensation.

## RESULTS

### Genome-wide CRISPRi screen reveals the ETC as a regulator of capsaicin-evoked death

Although many pathways have been suggested to account for TRPV1-mediated cell death, it is generally agreed that calcium influx is a required factor.^12,18^ Indeed, capsaicin-induced death of TRPV1^+^ HEK293T cells was abrogated by chelating or excluding extracellular Ca^2+^, demonstrating a similar requirement for Ca^2+^ influx in this heterologous system (Figure 1A). Moreover, both *Trpv1-EGFP*^+^ neurons (taken from *Tg(Trpv1-EGFP) MA208Gsat /Mmcd* mouse,^19^ see Figure S1) and TRPV1^+^ HEK293T cells showed similar morphological features of necrosis in response to capsaicin (Figure 1B and Video S1 and S2). This capsaicin-induced toxicity was not ameliorated by small molecule inhibitors of apoptosis, autophagy, or other distinct cell death modalities such as ferroptosis, necroptosis, pyroptosis and parthanatos (Figure S2),^20^ compelling us to take a more comprehensive and unbiased approach to identifying pathways involved in capsaicin-evoked excitotoxicity.

**Figure 1.**
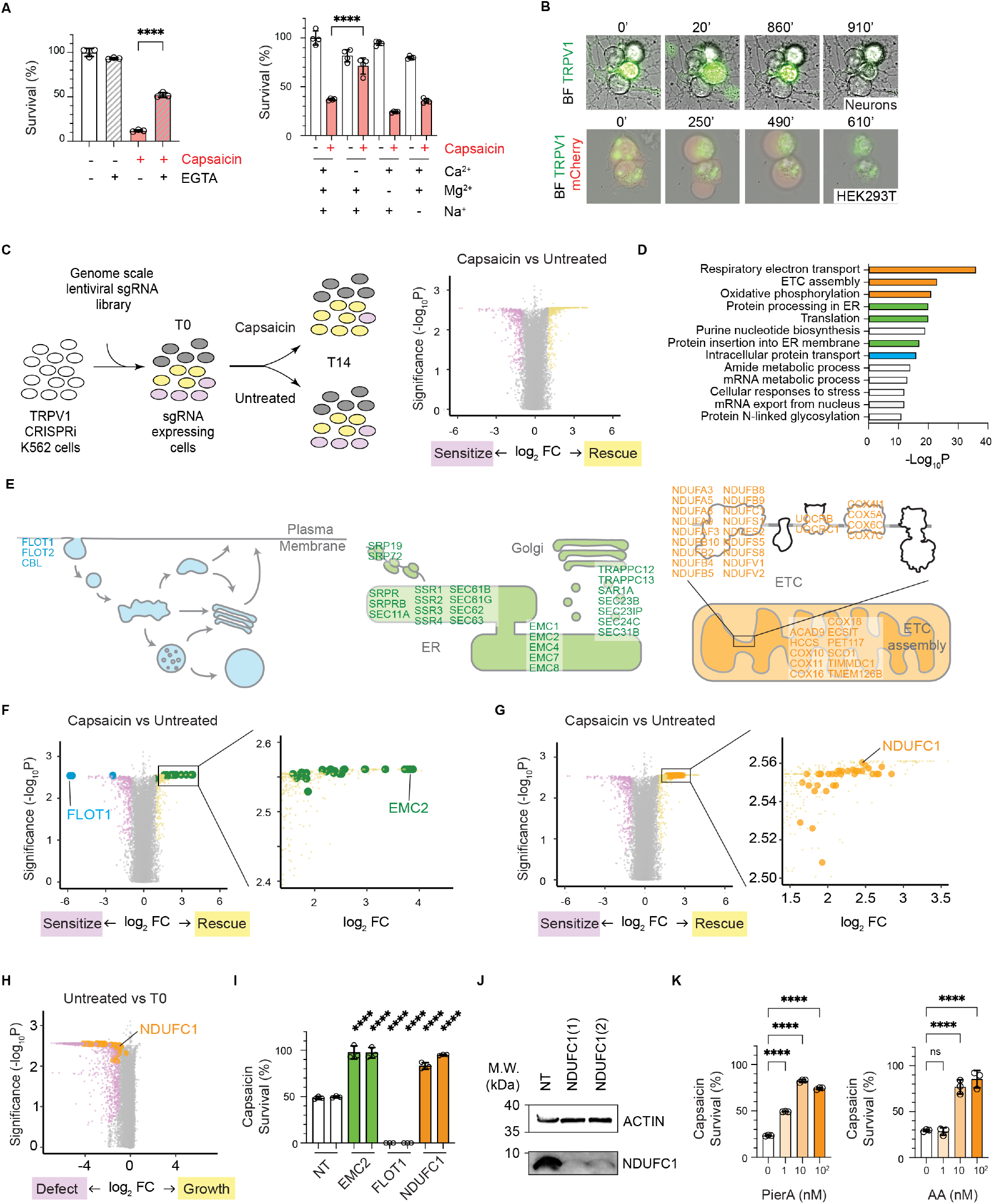
Genome-wide CRISPRi screen reveals the ETC as a regulator of capsaicin-evoked death. **(A)** Survival of TRPV1^+^ HEK293T cells following capsaicin or vehicle treatment under different ionic conditions: (left) Extracellular calcium was either chelated with EGTA; (right) indicated ions were dropped out of Ringer’s solutions that cells were in throughout the viability assay. **(B)** Representative images of DRG neurons from *Tg(Trpv1-EGFP)MA208Gsat/Mmcd* mice (top, Figure S1) and HEK293T cells transfected with plasmids containing GFP-TRPV1 and mCherry (bottom) from time-lapsed imaging following the addition of capsaicin in minutes above each image. TRPV1^+^ neurons and cells showed features of necrosis, including soma swelling and the eventual rupture of plasma membrane. **(C)** K562 cells engineered with TRPV1 and CRISPRi machinery were transduced with a genome-wide sgRNA viral library. Duplicated samples were collected before treatment (T0) and after 14 days of no or pulsated capsaicin (LD50) treatment and sequenced for the abundance of barcoded sgRNAs. Volcano plot shows the relative abundance of all targeted genes and non-targeting controls when comparing capsaicin (CAP) to untreated (UT) samples. **(D)** Most enriched Gene Ontology (GO) terms of CRISPRi screen hits of CAP compared to UT. **(E-G)** Strong hits in the endocytic (blue), membrane protein biogenesis (green) and ETC (orange) pathways are illustrated in cartoons and highlighted on volcano plots. **(H)** ETC hits from (G) is highlighted on the volcano plot that compares the relative abundance of sgRNAs from UT to T0. **(I)**Knocking down indicated genes with two sgRNAs leads to protection (EMC2, NDUFC1) or sensitization (FLOT1) against capsaicin (LD50) comparing to non-targeting controls (NT) clones. Two sgRNAs of each target gene and NT controls were used to make stable clones, and the capsaicin survival was normalized to vehicle control of each clone (vehicle controls are not shown here, see full figure in Figure S3). **(J)** Immunoblotting of lysates from NDUFC1 knockdown clones made from two different sgRNAs and a NT control. Actin was used as loading control. **(K)** Survival following capsaicin treatment was determined from cells pretreated with indicated concentrations of ETC inhibitors PierA or AA.For B,F,G, n 3 technical replicates representative of at least 3 independent experiments. Data are mean SD. One-way ANOVA Tukey’s or Dunnett’s multiple comparisons tests. **** P<0.0001.

Being able to recapitulate capsaicin-evoked death in an immortalized mammalian cell line enabled us to achieve this goal by conducting a genome-wide chemical genetic screen (Figure 1C).^11,17^ For this purpose, we engineered a TRPV1^+^ K562 clonal cell line expressing the required CRISPRi machinery (Figure S3A).^21^ These cells were transduced with a well-validated genome-scale single-guide RNA (sgRNA) library that targets 18905 genes with 5 sgRNAs/gene and 1895 non-targeting (NT) sgRNA controls.^22^ Samples were collected from two biological replicates at the outset of the experiment (T0) and from either untreated (UT) or capsaicin treated cells (CAP) at day 14, the experimental endpoint (Figure 1C and S3B). The effect of knocking down each gene on cellular survival was quantified by the relative abundance of targeting sgRNAs to NT controls for each gene.^21^ Using a 1% false discovery rate (FDR), we identified 1815 genes that regulate capsaicin-induced cell death by comparing UT and CAP sample groups and 2044 genes involved in cell growth by comparing T0 and UT sample groups (Table S1 and S2). Among gene hits that regulate capsaicin-induced cell death, 393 genes were regarded as “strong hits” with the absolute product of Log_2_FC and -Log_10_P > 4.

Among these “strong” candidate regulators, significant enrichment was seen in pathways that directly affect levels of TRPV1 on the cell surface by controlling membrane protein synthesis and intracellular protein transport, thereby validating the screen (Figure 1D and 1E). For example, knocking down a component of the ER membrane protein complex (EMC) led to reduced capsaicin toxicity by inhibiting the biogenesis of membrane proteins that include TRP channel family members (Figure 1E, 1F, and 1I).^23,24^ In contrast, knocking down flotillin (FLOT)-1, a protein that mediates endocytosis independent of clathrin and caveolin,^25,26^ exacerbated capsaicin-induced cell death. In addition to validating our screen, these ‘hits’ suggest that cell death is mediated through activation of TRPV1 at the cell surface. Consistent with this, we found that a cell-impermeant TRPV1 activator (the DkTx spider toxin)^27^ also confers cellular toxicity (Figure S3C).

The most enriched pathway identified in this screen corresponds to respiratory electron transport (ETC) (Figure 1D-1G, and Table S1). Specifically, components (25 genes) or assembly factors (11 genes) of the mitochondrial ETC strongly suppressed capsaicin-induced cell death (out of the 393 “strong hits”). Our finding is somewhat counterintuitive since disruption of the ETC and resulting metabolic consequences are usually deleterious to cell health.^7,28,29^ This conventional role of the ETC is also confirmed by our screen, as cells with hit sgRNAs of the ETC were more depleted at NT endpoint compared to T0 (Figure 1H). In contrast, we found that ETC disruption rescued capsaicin-evoked lethality (Figure 1G), indicating that ‘hits’ from the CRISPRi screen are revealing a unique regulatory strategy that balances resilience to excitotoxic insult against the cell’s overall metabolic needs.

To validate involvement of the ETC, we generated stable knock-down clones that targeted the mitochondrial complex I subunit *NDUFC1*, because it scored highly as a rescue hit against capsaicin-induced toxicity but had minimal effect on cell growth (Figure 1G and 1H). Indeed, knocking down *NDUFC1* protected against capsaicin-induced toxicity with minimal growth phenotype (Figure 1I, 1J and S3D). In contrast, knocking down *NDUFC1* in parental cells that do not express *TRPV1* showed no survival difference, ruling out any off-target effect of capsaicin treatment in NDUFC1-deficient cells (Figure S3E). To further corroborate this unexpected link between ETC suppression and excitotoxicity, we tested the effect of ETC inhibition on capsaicin-induced cell death using a collection of well-defined small molecule inhibitors. Pretreatment with the complex I inhibitor, piericidin A (PierA) (1-10 nM), suppressed toxicity by capsaicin (Figure 1K). Similar protective effects were observed with low concentrations of the complex III inhibitor, antimycin A (AA), and two other complex I inhibitors, rotenone and phenformin (10 nM, 10 nM and 10 µM respectively) (Figure 1K, S3F, and S3G). High concentrations of these inhibitors can promote apoptosis, but as previously reported,^30^ only mild decreases of growth rates were observed at the relatively low concentrations that were protective against capsaicin lethality.

Because suppressed ETC activity, and thus oxidative phosphorylation, protects cells from capsaicin-induced toxicity, we next asked whether promoting aerobic respiration would exacerbate toxicity. Indeed, we found that cells were sensitized to capsaicin-induced death when cellular respiration was increased by reducing glycolysis (Figure S3H). The shift of metabolism from glycolysis to oxidative phosphorylation is mediated by the loss of the mRNA splicing factor, LUC7L2,^31^ which was also a sensitizing hit in our CRISPRi screen (Table S1).

By what mechanism might ETC inhibition protect cells from capsaicin-evoked death? As described below, we have identified two main pathways that rescue TRPV1-mediated cell death.

### Metabolic regulation of TRPV1 endocytosis

Change in metabolic state may affect TRPV1 status;^32^ we therefore asked whether ETC suppression imparts protection by inhibiting receptor expression, localization, or function. We found that *NDUFC1* knockdown or treatment with PierA led to a partial reduction of capsaicin-evoked Ca^2+^ influx (Figure 2A and 2B). To investigate the underlying mechanism, we utilized *FLOT1* knockdown cells that are defective at endocytosing TRPV1 (Figure 1F and 1I). We found that PierA also reduced capsaicin-induced Ca^2+^ influx in *FLOT1* knockdown cells (Figure 2C), suggesting that its effect on reducing Ca^2+^ influx is mediated by enhancing TRPV1 endocytosis.

**Figure 2.**
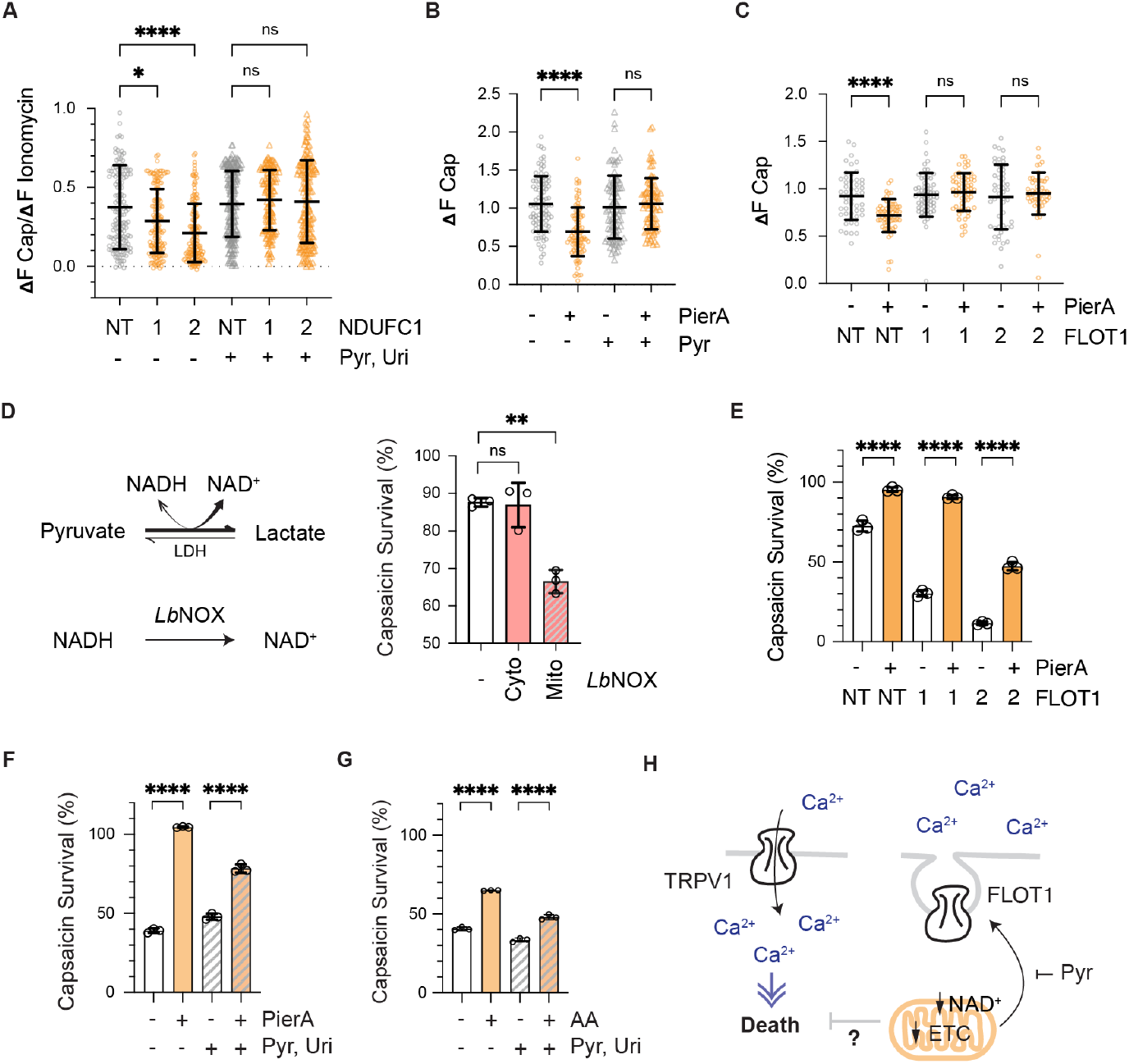
Metabolic regulation of TRPV1 endocytosis. **(A)**Calcium response was measured as change in FURA-2 AM ratio in response to capsaicin (33nM) as a fraction of maximum response to 10 µM ionomycin in TRPV1^+^ K562 knockdown clones of *NDUFC1* (using 2 different sgRNA) and a non-targeting (NT) control. **(B**,**C)** Maximum calcium response to 0.4 µM capsaicin following the indicated pretreatment using K562 parental (B), NT, or *FLOT1* knockdown clones generated by 2 sgRNAs as indicated (C). **(D)** Pyruvate accepts electrons from NADH to produce lactate and NAD^+^, a coupled redox reaction catalyzed by lactate dehydrogenase (LDH) that supports the proliferation of ETC-deficient cells. Oxidation of NADH to NAD^+^ can be carried out by *Lb*NOX. Viability of K562 stable lines that expressed empty vector, cyto, or mito*Lb*NOX after capsaicin treatment and normalized to vehicle control.**(E-G)** Following indicated pretreatments, viability of K562 cells was determined after treatment with capsaicin and normalized to vehicle controls of each pretreatment and genetic conditions. **(H)** Schematic summary: A fraction of ETC rescue is mediated through FLOT1 dependent TRPV1 endocytosis, which can be negated by increased pyruvate (and thus NAD^+^/NADH). An alternative rescue mechanism independent of endocytosis is also unmasked. Data are mean SD * P<0.05, ^**^ P<0.01, ^****^ P<0.0001. **(A-C)**: Kruskal-Wallis test with Dunn’s multiple comparisons test. n = (A) 150; (B) 90; or (C) 60 cells from 3 independent experiments. **(D-G)**: One-way ANOVA with Dunnett’s (D) or Tukey’s (E-G) multiple comparisons test. n = 3 technical replicates, representative of at least 3 independent experiments.

In another line of inquiry, we used pyruvate and uridine to uncouple downstream effects of ETC suppression. Addition of pyruvate and uridine to the culture medium is known to rescue proliferation of cells that lack a functional ETC.^33-35^ We found that the addition of pyruvate, in particular, negated the decrease of capsaicin-evoked Ca^2+^ influx caused by *NDUFC1* knockdown or PierA treatment (Figure 2A and 2B), demonstrating its role in regulating TRPV1 trafficking via FLOT1, a regulatory pathway that may also pertain to other ion channels, transporters, or receptors.^25,26,36,37^

Because the main role of pyruvate when added to ETC deficient cells is to regenerate NAD^+^,^35^ we asked whether direct manipulation of [NAD^+^]/[NADH] ratio can affect capsaicin lethality. Using a NADH oxidase from *Lactobacillus brevis* (*Lb*NOX),^38^ we found that capsaicin-induced toxicity is exacerbated when *Lb*NOX is targeted to the mitochondria (mito*Lb*NOX) but not the cytoplasm (cyto*Lb*NOX) (Figure 2D). As mito*Lb*NOX was reported to double cellular [NAD^+^]/[NADH] whereas cyto*Lb*NOX did not result in significant changes,^38^ our data suggest that increasing [NAD^+^]/[NADH], especially in the mitochondria, exacerbates capsaicin-induced toxicity. This is consistent with the protection towards capsaicin-induced lethality afforded by suppressing ETC function and its effect on lowering [NAD^+^]/[NADH].^35^ Moreover, our CRISPRi screen identified *SLC25A51*, the mammalian mitochondrial NAD^+^ transporter,^39^ as a strong protective hit (Table S1) agreeing with the protective effect of lowering [NAD^+^]/[NADH] in the mitochondria. Taken together, these data suggest that decreased [NAD^+^]/[NADH] contributes to the rescue effect of ETC suppression by regulating endocytosis, a process that can be overridden by pyruvate addition.

The above results support a trafficking-dependent pathway through which ETC suppression rescues capsaicin lethality. Interestingly, PierA or AA treatment still provided significant protection in *FLOT1*-deficient or pyruvate supplemented cells (Figure 2E-2G and S3I-S3K), revealing the presence of another protective mechanism orthogonal to endocytic regulation (Figure 2H). Going forward, we included pyruvate and uridine in our experiments to isolate and characterize this trafficking-independent mechanisms of rescue.

### Capsaicin and ETC dependent transcriptional changes

To probe the endocytosis-independent component of ETC rescue, we took an unbiased transcriptomic-based approach comparing cellular responses to capsaicin (with pyruvate and uridine supplementation) in the absence or presence of PierA. We reasoned that if a factor mediates capsaicin-induced toxicity, then its expression could be changed upon capsaicin treatment and this change should be dampened by PierA. Pair-wise analysis between vehicle and capsaicin treated samples showed prominent transcriptional changes in “capsaicin responsive genes” after 4 hours of treatment (Figure 3A and Table S3). Gene ontology analyses showed that transcription factors were the dominant class of differentially expressed genes, of which many are known to respond to calcium and / or various forms of cellular stress. Importantly, PierA pretreatment dampened transcriptional changes in these capsaicin responsive genes (Figure 3B).

**Figure 3.**
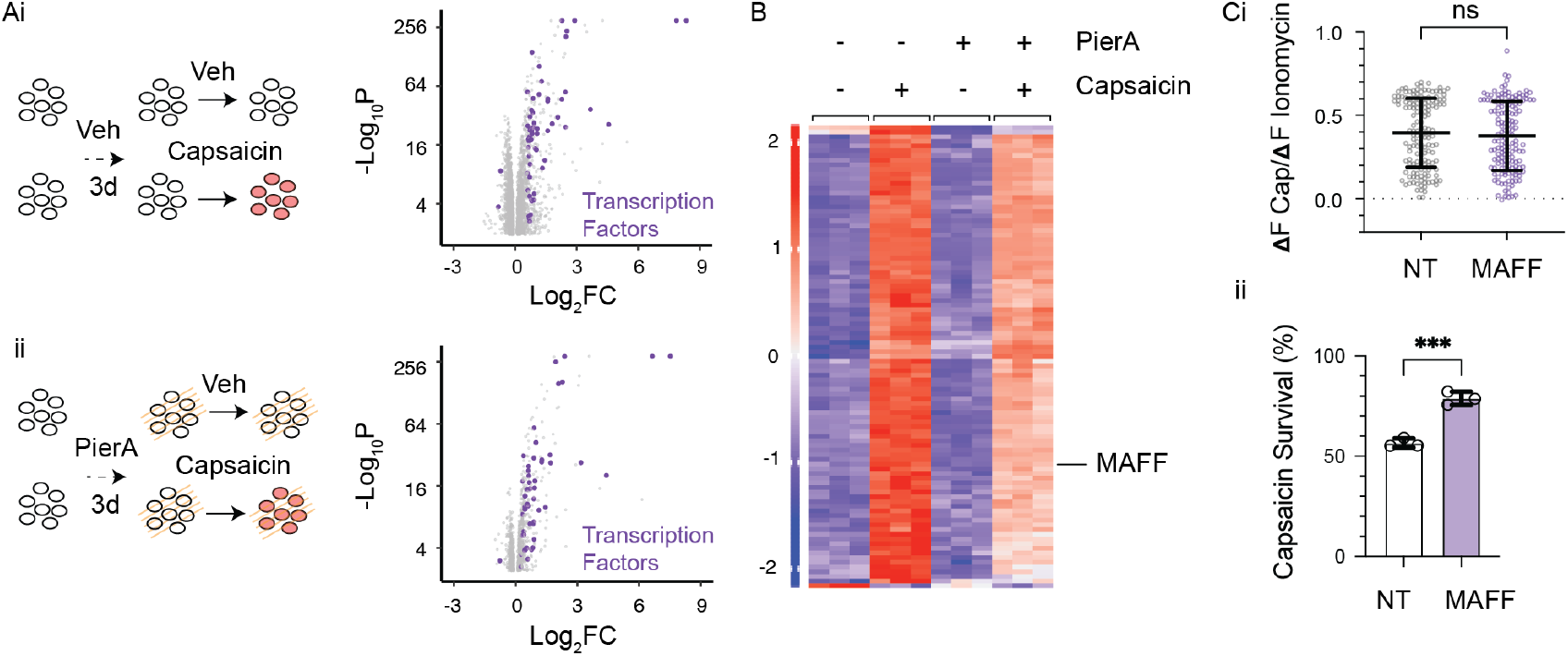
Capsaicin and ETC dependent transcriptional changes. **(A)** Schematic of RNAseq experiment. RNAseq analyses were performed on 3 replicates of each condition. Differentially expressed genes of capsaicin versus vehicle treatments following (i) vehicle or (ii) PierA pretreatment were plotted. Transcription factors are colored purple.**(B)** Relative expression of top 100 significantly different genes in response to capsaicin **(A**i) was plotted based on the Z score of normalized counts (rlog transformation of the DEseq2 package). **(C)** K562 cells that stably expressed NT or *MAFF*-targeting sgRNAs were tested for their calcium response to capsaicin (i) (assay see Figure 2A) and viability after capsaicin treatment was quantified and normalized to vehicle (ii). Data are meanSD. **(C**i): Mann Whitney test. n = 150 cells from 3 independent expriments; **(C**ii): unpaired 2-tailed t-test. n = 3 technical replicates, representative of 3 independent experiments. *** P<0.001.

Cross-referencing capsaicin-responsive transcription factors with the CRISPRi screen hits revealed one candidate, *MAFF* (Table S1 and S3), which encodes a small DNA binding protein, MafF. MafF is recruited to antioxidant response elements during oxidative stress, and either activates or represses transcription depending on its binding partners.^40^ *MAFF* is upregulated upon capsaicin treatment and its upregulation is dampened by PierA (Figure 3B). Knocking down *MAFF* mitigated capsaicin-induced cell death without affecting Ca^2+^ influx (Figure 3C). Together, these data suggest that capsaicin-induced upregulation of *MAFF* accounts for some component of toxicity. Although MafF can also bind to calcium-responsive elements, this profiling was carried out in the presence of pyruvate to minimize the endocytosis-dependent component of rescue (Figure 2B). We therefore focused on elucidating the relationship between ETC suppression and oxidative stress in capsaicin-evoked excitotoxicity.

### ETC suppression boosts resilience by lowering oxidative stress

As electrons pass down the ETC, they occasionally leak prematurely and generate reactive oxygen species (ROS) such as superoxide anions (O_2_^•-^) and hydrogen peroxide (H_2_O_2_).^41,42^ We first assessed changes in mitochondrial ROS generation upon capsaicin treatment with mitoSOX, a mitochondrially targeted fluorescent indicator of superoxide. Using flow cytometry, capsaicin exposure increased the percentage of clonal TRPV1^+^ HEK293T cells showing a high mitoSOX signal; no such shift was observed in parental lines not expressing *TRPV1* (Figure S4A). To investigate this phenomenon further, we targeted a genetically encoded ratiometric H_2_O_2_ indicator HyPer7 to the mitochondria matrix of TRPV1^+^ HEK293T cells, which revealed a steady increase of mitochondrial H_2_O_2_ in response to capsaicin that plateaued within 30 min (Figure 4A).^43^ Moreover, omitting extracellular Ca^2+^ during the capsaicin treatment prevented the increase of mitochondrial HyPer7 and mitoSOX signals, supporting the critical role of Ca^2+^ in ROS generation (Figure 4A, S4B, and S4K).^2,42^

**Figure 4.**
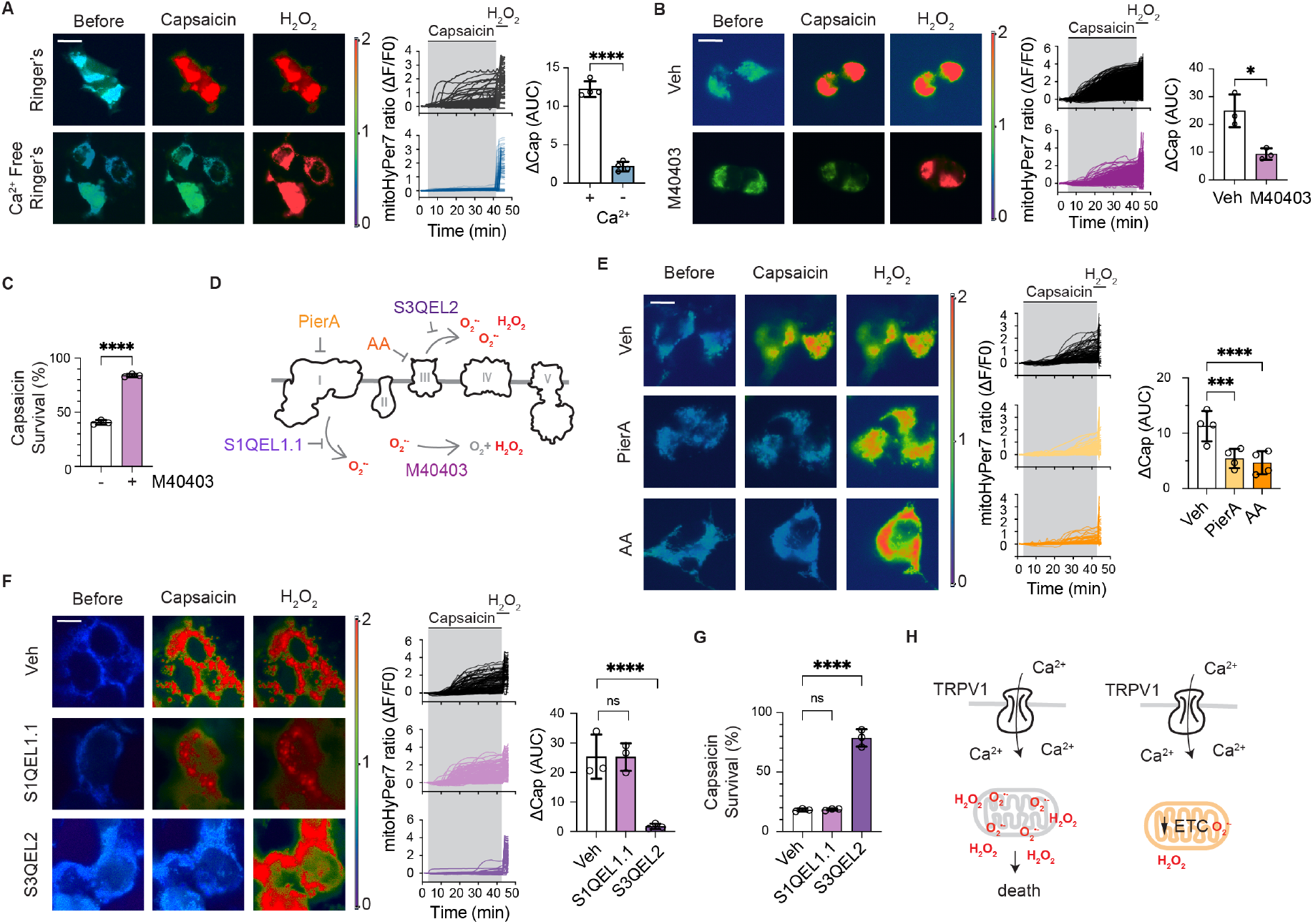
ETC suppression boosts resilience by lowering oxidative stress. **(A**,**B**,**E**,**F)** Mitochondrial ROS generation kinetics in TRPV1^+^ HEK293T cells were revealed by time-lapsed microscopy of mitoHyPer7. Capsaicin (20 µM) was added at 3 min and H_2_O_2_ (200 µM) at 42 min to elicit maximum response under indicated conditions. Representative images of the same cells at T0 (before), T42 (max capsaicin response), and T45 (max H_2_O_2_ response) are shown in pseudo color of HyPer7 ratio (ex480/ex380). Cell response kinetics are shown as changes of HyPer7 ratio (ΔF/F0) over time in a representative experiment, and averaged capsaicin response measured by area under curve (AUC) of experimental replicates are shown. **(A)** Capsaicin-evoked HyPer7 response in cells imaged in regular or calcium free Ringer’s. **(B**,**C)** HyPer7 response (B) and viability (C) of cells pretreated with a SOD memetics M40403. **(D)** Schematics of small molecule inhibitors used in this Figure. Pier A and AA are inhibitors of complex I and III respectively. S1QEL1.1 and S3QEL2 block ROS production via sites I_Q_ and III_QO_ on complex I and III respectively. M40403 detoxify O^•-^ to H_2_O_2_.**(E)** Cells pretreated with ETC inhibitors PierA or AA (with pyruvate and uridine) were examined for their capsaicin-evoked HyPer7 response. **(F**,**G)** Cells pretreated with vehicle, S1QEL1.1, or S3QEL2 were treated with capsaicin and their HyPer7 response (F) and survival (G) were examined. **(H)** Schematic summary: capsaicin elicits Ca^2+^ influx by activating TRPV1, which leads to mitochondrial ROS generation and cell death. ETC suppression decreases capsaicin-evoked ROS generation, which is orthogonal to its endocytic regulation. Data are meanSD. **(A**,**B**,**E**,**F)**: two-way ANOVA (A,B) with Dunnett’s multiple comparisons (E,F). N = (A, E) 4; or (B,F) 3 as number of independent experiments (see Figure S4K). **(C)**: 2-tailed unpaired t-test. **(G):** one-way ANOVA with Dunnett’s multiple comparisons test. (C,G) n = 3 technical replicates, representative of at least 3 independent repeats. *P<0.05, *** P<0.001, **** P<0.0001.

The mitochondrial calcium uniporter (MCU) has been proposed to mediate Ca^2+^ clearance at the synapse^44^ as well as glutamate toxicity.^2^ By constructing a TRPV1^+^ MCU knockout cell line,^45^ we found that MCU is dispensable for capsaicin-induced cell death (Figure S4C). Consistent with this, sgRNAs that target MCU components did not emerge as hits in our CRISPRi screen (Table S1). Moreover, capsaicin-induced Ca^2+^-dependent increase of ROS was also evident in cells that lacked MCU (Figure S4D), consistent with its dispensable role in capsaicin-evoked cell death. These data suggest that either Ca^2+^ flux across the mitochondrial inner membrane is not critical for capsaicin-evoked ROS generation and lethality, or that calcium enters mitochondria to generate ROS through a MCU-independent mechanism, such as via the Ca/H exchanger LETM1 or Ca/Na exchanger NCLX in the mitochondrial inner membrane^46^ in our model system.

To test whether capsaicin-evoked ROS production leads to cell death, we pretreated cells with M40403, a superoxide dismutase (SOD) memetic, prior to capsaicin treatment. M40403 dampened the capsaicin-induced increase in the HyPer7 and mitoSOX signals and rescued capsaicin-induced cell death without altering calcium dynamics (Figure 4B-4D, S4E, S4F, and S4K). The moderate decrease of the HyPer7 response in M40403 treated cells is consistent with HyPer7 being most sensitive to H_2_O_2_,^43^ but which may also respond to high concentration of O_2_^•-^. Pretreatment with mitoTEMPO, another mitochondria-targeted SOD memetic, also significantly ameliorated capsaicin-induced cell death (Figure S4G). The effective protection of these SOD memetics against capsaicin-induced cell death is consistent with the identification of *SOD1* as a sensitizing hit from the CRISPRi screen (Table S1). These data suggest that capsaicin damages and kills cells by increasing ROS generation.

We next asked if ETC suppression mitigates capsaicin-evoked ROS generation. Indeed, pretreating cells with PierA or AA (supplemented with pyruvate and uridine) diminished the capsaicin-induced increase of HyPer7 and mitoSOX signals and rescued survival (Figure 4E, S4H, S4I, and S4K). Although AA is known to evoke ROS generation at high concentrations, we did not notice a significant change of baseline HyPer7 ratio before the addition of capsaicin in cells pretreated with low nano molar AA in our experiments. Our data show that mild ETC suppression can mitigate capsaicin-evoked enhancement of oxidative stress and ameliorate its downstream deleterious effect.

To further understand capsaicin-evoked ROS generation, we tested the effect of S1QEL1.1 and S3QEL2, which specifically suppress O_2_^•-^ or H_2_O_2_ generation from two major ETC sites without affecting ETC function.^42,47,48^ Interestingly, capsaicin-induced ROS generation and cell death were negated by S3QEL2 but not S1QEL1.1, suggesting ROS is produced by site III_QO_ on complex III (Figure 4F, 4G, S4J, and S4K).

As site III_QO_ faces the inner membrane space, it is consistent with the identification of the inner membrane / cytosolically localized SOD1 but not matrix localized SOD2 as a sensitizing hit from our CRISPRi screen. Taken together, our data suggest that suppression of Ca^2+^-induced mitochondrial ROS represents a major mechanism for ETC protection against capsaicin-evoked death (Figure 4H).

### Endogenous ETC level inversely correlates with sensory neuron resilience

Based on the role of the ETC in modulating capsaicin-induced lethality in heterologous immortalized cells, we investigated the expression of ETC in adult mouse sensory nociceptors. Interestingly, reanalysis of a published RNAseq dataset (GEO: GSE131230) shows that ETC components are expressed at lower levels in TRPV1^+^ neurons compared to other sensory neuron subtypes (Figure 5A).^49^ Moreover, immunohistochemical staining of sensory ganglia with antibodies against two ETC components confirmed their reduced expression in TRPV1^+^ neurons (Figure 5B and S5A). Consistent with this, TRPV1^+^ sensory neurons were more resilient to apoptosis induced by high concentrations of ETC inhibitors compared to the TRPV1^-^ population, suggesting less relative dependency on the ETC (Figure 5C), as recently described for oocytes in dormancy.^50^

**Figure 5.**
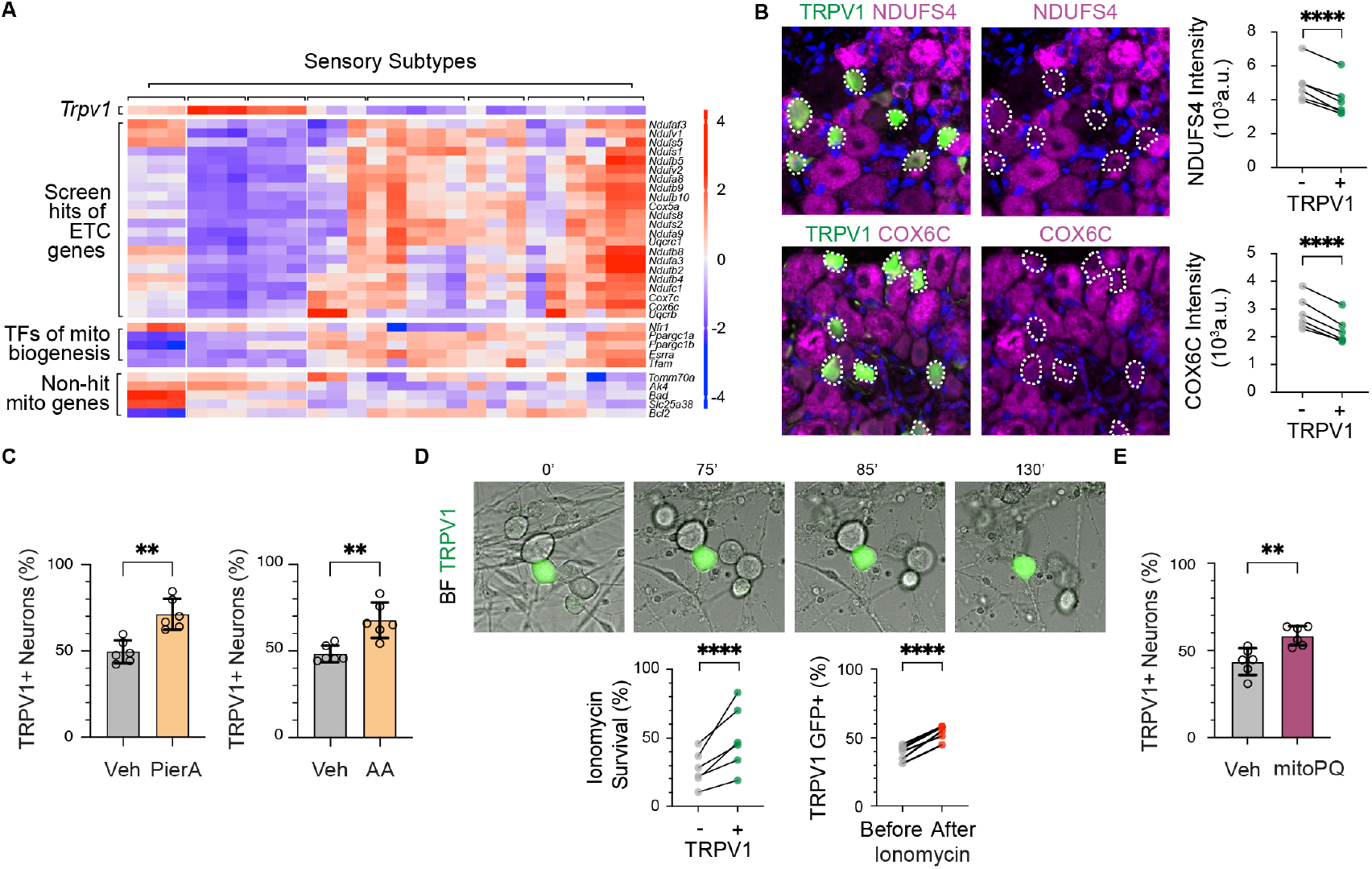
Endogenous ETC level inversely correlates with sensory neuron resilience. **(A)** Normalized expression level (rlog) among different sensory subgroups were plotted as Z scores for genes encoding TRPV1, ETC components identified by the CRISPRi screen as regulators, transcription factors of ETC biogenesis, and mitochondrial genes that did not mediate capsaicin-induced toxicity in the screen as controls. Heatmap was generated using a published dataset (GEO: GSE131230). **(B)** Representative images of DRG sections from TRPV1-GFP mice (Figure S1) immunolabeled with antibodies that target ETC components (NDUFS4 or COX6C) and GFP. Labeling intensities of ETC components in TRPV1(GFP)^+^ were quantified and compared TRPV1(GFP)^-^ neurons. **(C)** After treating primary DRG culture with high levels of ETC inhibitors PierA or AA (with pyruvate and uridine) and a recovery period, percentage of TRPV1^+^ neurons were calculated from cells that showed calcium response to capsaicin and high extracellular K^+^ (latter was used to reveal all neurons) by FURA-2AM. **(D)** Representative images of DRG neurons taken from TRPV1-GFP mice (Figure S1) undergoing necrosis after ionomycin addition in minutes indicated above each image. Survival of TRPV1(GFP)^+^ or TRPV1(GFP)^-^ after ionomycin treatment and percentage of TRPV1(GFP)^+^ before and after ionomycin treatment were quantified from microscopy videos. **(E)** After treating cells with mitoPQ, percentage of TRPV1^+^ neurons were determined as described in (C). **(B**,**D)** Data are mean per animal and lines connect the same animal. N = 6 animals. Two-way ANOVA (data of each animal see Figure S5). **(C**,**E)** Data are mean per animal with SD. N = 6 animals, calculated from at least 3 technical replicates per animal. Unpaired 2-tailed t-test. ^**^ P<0.01; ^****^ P<0.0001.

We next asked whether TRPV1^+^ neurons are also more resistant to excitotoxic or oxidative stress due to their lower expression of ETC components. Because capsaicin is selectively toxic to TRPV1^+^ neurons, we used ionomycin, a bacterial calcium ionophore that facilitates the diffusion of calcium ions across cell membranes, to evoke sustained calcium influx and excitotoxicity to all cells. Time-lapse imaging showed that 10 µM ionomycin with 10 mM extracellular Ca^2+^ induced cellular swelling and necrosis in primary dorsal root ganglia (DRG) culture (Figure 5D, Video S3). Interestingly, TRPV1^+^ sensory neurons consistently fared better under this excitotoxic challenge compared to the TRPV1^-^ neurons (Figure 5D and S5B). We also showed that TRPV1^+^ neurons were more resilient to oxidative stress by exposing DRG cultures to the mitochondrial ROS generator mitoParaquat (mitoPQ) (Figure 5E).^51^

### ETC regulates sensory neuron resilience

The above data suggest that relative sensitivity of sensory neurons to excitotoxic and oxidative stress is determined by their level of ETC expression as observed in our heterologous cell culture experiments (Figure 1I and 1K). To test this hypothesis, we manipulated ETC expression in sensory neurons using orthogonal genetic strategies. To achieve loss-of-function, we conditionally ablated *Ndufs4* or *Uqcrq* (complex I or III component respectively)^52,53^ from TRPV1 lineage neurons (*Trpv1*^*Cre*^),^54^ which reduced their viability while also imparting protection against capsaicin (Figure 6A-6C). This dual effect is consistent with both the essential role of the ETC for neuronal survival past 7 weeks of age ^52^ and the protective effect of reduced ETC expression that we observe when cells are challenged with capsaicin.

**Figure 6.**
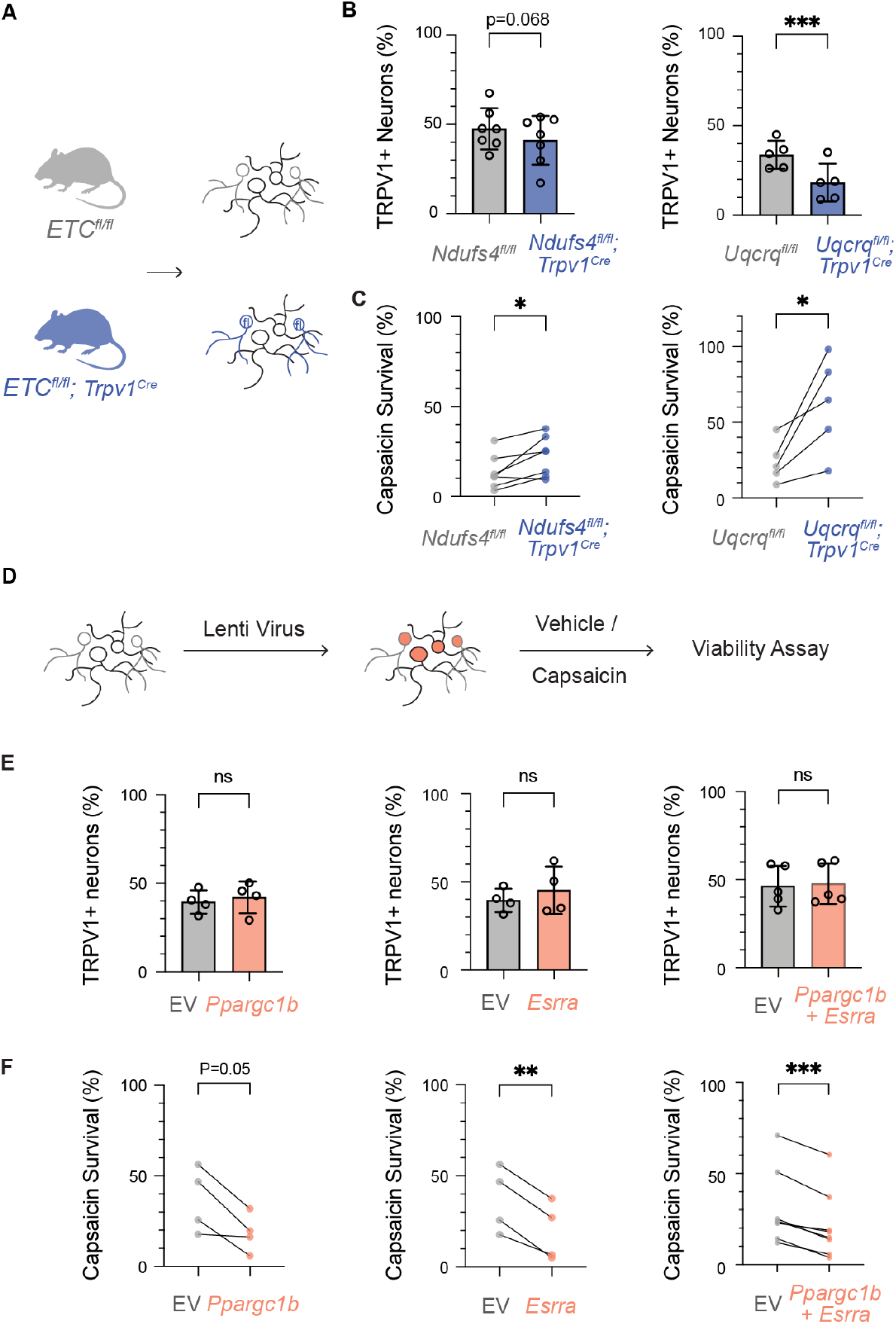
ETC regulates sensory neuron resilience. **(A)** Schematics illustrates experimental design of (B,C). **(B**,**C)** DRG cultures from littermates of the indicated genotypes were treated with vehicle or capsaicin. Percentage of TRPV1^+^ neurons in vehicle treated conditions and the percentage of TRPV1^+^ neurons survived capsaicin treatment were calculated from the number of TRPV1^+^ and total neurons at end points using FURA-2AM assays (see Figure S6 and Methods). N= 7 or 5 pairs of animals for *Ndufs4* or *Uqcrq* flox, calculated from least 3 technical replicates per animal. **(D)** Schematics illustrates experimental design of (E,F). **(E-F)** Primary DRG neurons were transduced with lentivirus carrying empty vector (EV) or indicated genes and treated with vehicle or capsaicin. Similar assay as (B,C). N 4 biological replicates, calculated from at least 3 technical replicates. Data are mean SD. **(B**,**E)** Unpaired 2-tailed t-test. **(C**,**F)** Paired 2-tailed t-tests where each animal / biological replicate were paired due to baseline variation; ^*^P<0.05; ^**^P<0.01; ^***^P<0.001; ^****^P<0.0001.

To achieve gain-of-function, we ectopically expressed *Ppargc1b* and *Esrra* to boost ETC biogenesis. These genes encode transcription co-factors PGC1β and ERRa, which were chosen over other known factors because they are both screen hits and differentially expressed by the subpopulation of TRPV1^+^ sensory neurons (Figure 5A and Table S1).^55,56^ Indeed, sensory neurons that overexpressed *Ppargc1b, Esrra*, or a combination were significantly more sensitive to capsaicin-induced lethality compared to those transduced with empty vector (EV) controls, while the percentage of TRPV1^+^ neurons remained unchanged (Figure 6D-6F). Therefore, increasing ETC biogenesis weakened the resilience of TRPV1^+^ nociceptors to excitotoxicity.

## DISCUSSION

Neurons are post-mitotic and thus the consequences of damage or death are dire compared to proliferative cells. Primary afferent sensory neurons can regenerate their peripheral terminals, a process that is dependent on local calcium transients as well as energy provided by mitochondria. ^57-60^ Interestingly, a recent study showed that nociceptors regenerate their peripheral terminals more robustly than tactile nerve fibers.^61^ Of course, nerve terminal regeneration is predicated on survival of the soma, the resilience of which has not, to our knowledge, been investigated using a comprehensive chemical genetic screen. Our genetic and pharmacologic dissections support a role for ETC tuning in determining sensory neuron resilience. While our investigation began by using capsaicin as a model, we also show that this conclusion pertains to other forms of excitotoxic and oxidative insults (Figure 5D and 5E).

### Metabolic tuning as a regulatory mechanism

Given its critical role in ATP production and NAD+ regeneration, impairing ETC activity generally has negative consequences for cellular health, as we indeed observed with both proliferating cell models and neurons (Figure 1H and 6A).^52^ However, we show that maintaining low ETC activity can be beneficial in the face of excitoxicity, revealing interesting parallels to other physiologically challenging scenarios. For example, knocking down expression of ETC components delays aging in many model organisms.^62-64^ Another more extreme example was recently described for early oocytes, which eliminate their complex I altogether to avoid ROS accumulation during dormancy, when energy expenditure is presumably minimal.50 Furthermore, the idea that ETC activity plays a regulatory role is also consistent with recent observations that species-specific developmental rates are determined by mitochondrial function and metabolic rate.^65,66^ While the abovementioned studies focused on steady state ROS production, our study highlights the role of ETC suppression in regulating calcium-dependent ROS generation, which is especially relevant to excitoxicity and pathological conditions that accompanied painful neuropathies.

Moreover, we uncovered an unexpected downstream effect of ETC regulation on intracellular trafficking. Specifically, our screen highlights a new metabolic regulation of excitotoxicity via the flotillin-dependent endocytic pathway. Flotillins are known to mediate the assembly and endocytosis of large protein complexes in sphingolipid/cholesterol rich microdomains on the plasma membrane^.25,26^ As many ion channels, including Kv2.1, have been reported to cluster and/or localize to lipid rafts,^37,67,68^ this pathway may act broadly, beyond TRPV1 in primary afferent neurons. Further analysis of this pathway may provide insights as to why dysregulated metabolic conditions, such as obesity or diabetes, are associated with chronic pain syndromes.

### Differential ETC expression tuned to the heterogenous needs of different sensory subtypes

We show that a major class of nociceptors maintains lower ETC expression compared to other sensory subtypes, granting them greater resilience against excitotoxic and oxidative stress (Figure 5). We speculate that several factors, including ion channel physiology, cellular properties, and pathogen tropism, may be driving this difference among sensory neuron subtypes.

The TRPV1 ion channel has biophysical properties that may uniquely dispose cells to excitotoxicity. It is highly Ca^2+^ permeable compared to other excitatory ion channels in peripheral sensory neurons (such as ASICs, P2X, or other TRP channels^15,69^) or more slowly desensitizing through a process that likely involves endocytosis^70^ thus conferring a higher risk of calcium overload. Additionally, inflammatory mediators potentiate TRPV1 activation, further increasing the risk of excitotoxicity under maladaptive conditions.^13,14^

Interestingly, many nociceptive neurons also express *Trpa1* channels,^49^ a receptor for a wide variety of reactive electrophilic irritants, including endogenous products of oxidative stress such as 4-hydroxynonenal or 15-deoxy-Delta(12,14)-prostaglandin J(2).^71-74^ Low ETC expression reduces ROS generation, which may protect the cell from chronic activation of TRPA1 or other ROS-sensitive ion channels to further limit toxicity. On the other hand, nociceptors may have lower energic demands as they usually fire non-repetitively or at much lower frequency compared to other subtypes.^49^ For example, Aβ low threshold fibers that specialize in detecting high-frequency vibratory stimuli can fire over 500 Hz.^75^ As calcium influx during an action potential is smaller and more transient compared to that associated with TRPV1 activation, excitoxicity during high frequency firing is less of a risk and likely outweighed by the higher ETC activity needed to support the energetic needs of these light touch neurons. Thus, different sensory neuron subgroups may adjust their metabolic states to suit their diverse functional roles.

Finally, tropism of pathogens can contribute to the selective pressure that drives the differential requirement of resilience. For example, *Staphylococcus aureus*, including antibiotic-resistant strains, secrets pore-forming toxins that preferentially attack TRPV1^+^ neurons to induce calcium influx, action potentials, and pain sensation.^76,77^ Resilience against such bacterially-evoked calcium overload is desirable as TRPV1^+^ neurons also orchestrate protective immunity via the release of neuropeptides such as CGRP.^14,78^

### Mechanisms of capsaicin-induced excitotoxicity

Our characterization of TRPV1-mediated excitotoxicity shows that, like glutamate toxicity, calcium overload and oxidative stress are key steps leading to cell death. We found that ROS produced by site III_QO_ of the mitochondrial ETC contributes to capsaicin-induced cell death (Figure 4F and 4G). Although we do not pinpoint the exact ROS species involved, the effective lethality rescue using SOD memetics (Figure 4C and S4G) and our identification of SOD1 as a sensitizing hit suggests that O_2_^•-^ may be the more relevant culprit. As SODs catalyze the conversion of O_2_^•-^ to the less reactive H_2_O_2_, it is consistent to find overlapping hits between our screen and other CRIPSR and shRNA screens that used exogenously added H_2_O_2_ to induce oxidative stress.^79^ Conversely, genes or pathways unique to our screen may play a role in the detoxification of the more deleterious O_2_^•-^ and warrant further investigation in the future.

The case is perhaps more complex for glutamate-evoked toxicity, where oxidative stress can originate from several sources, including nitric oxide synthase (NOS)^80^ and NADPH oxidases (NOXs)^81^. In this case, a key cell death executioner downstream of oxidative stress is poly(ADP-ribose) polymerase (PARP)-1,^1,2,82,83^ which initiates a cell death modality called “parthanatos” by catalyzing polyADP-ribosylation (PARylation), where the main cellular donor of ADP-ribose moiety is NAD^+^. Indeed, disruption of NAD^+^ metabolism plays a key role in axon degeneration of PNS neurons, as exemplified by the studies of Wallerian degeneration slow (*Wld*^*S*^) and SARM1.^84^ Although we found that increasing NAD^+^/NADH ratio in mitochondria sensitizes cells towards capsaicin-induced toxicity (Figure 2H), pretreatment with a potent PARP-1 inhibitor did not protect against capsaicin-evoked cell death (Figure S3B), suggesting involvement of other pathways. Furthermore, small molecule inhibitors of lipid peroxidation and ferroptosis failed to rescue capsaicin-evoked toxicity (Figure S3C), nor did our CRISPRi screen reveal known regulators of the parthanatos or ferroptosis pathways, suggesting that another oxidative stress evoked cell death pathway(s) acts as executioner.

Although the exact pathway remains to be elucidated, we identified the small transcription factor, MafF, as a regulator of this cell death modality. Going forward, it will be important to understand how MafF senses ROS and regulates cell death. Recently, another nuclear ROS sensor, CHK1, was reported to dampen ROS generation and mediate resistance to platinum-based anticancer agents by blocking translation in mitochondria via SSBP1.^85^ As site IQ but not III_QO_ was responsible for ROS generation in their system, it is tantalizing to ask if CHK1 and MafF can mount site-specific responses. Interestingly, expression of MafF in brain endothelial cells was recently shown to be age-dependent.^86^ As aging is a major risk factor of painful neuropathy, it will be interesting to ask if changes in MafF expression contribute to decreased resilience during aging.

In summary, we propose that TRPV1^+^ sensory neurons establish a metabolic ‘sweet spot’ in which reducing ETC expression protects them from excitotoxicity while accommodating energetic demands of maintaining electrical excitability over long and arborized processes that innervate distant receptive fields. This balancing act between environmental resilience and metabolic sufficiency may limit the extent to which these neurons are protected against chronic or severe challenges. Therefore, it will be interesting to determine whether and to what extent modulating the ETC can be beneficial in pathological situations that lead to painful neuropathy, including disorders such as diabetes that alter metabolic homeostasis.^7^

## Acknowledgments

We thank A. Akopian for providing the *Tg(Trpv1-EGFP)MA208Gsat/Mmcd* mouse line, J. Weissman and M. Kampmann for CRISPRi cell lines and reagents. We thank A. Samelson, J. Nunez, M. Shurtleff, and M. Jost for advice on the CRISPRi screen, N Ingolia for providing computational resources for sequencing analyses, and Y Kirichok for WT and KO MCU cell lines. We thank Y Kirichok, K Birsoy, J Nikkanen, K Yackle, R Nicoll and members of the Julius lab for discussion and critical reading of the manuscript. We appreciate technical support and advice from UCSF core facilities: S Elmes (UCSF LCA NIH Cancer Center Support Grant P30CA082103), D Larsen, K Herrington and SY Kim (UCSF CALM), and E Chow and D Martinez (UCSF CAT). This work was supported by Warren Alpert Distinguished Scholar Fellowship (L.Y.); Larry L. Hillblom Foundation Fellowship (L.Y.); University of California San Francisco Cardiovascular Research Institute T32 (L.Y.); National Institute of Health grant P01AG049665 (N.S.C); National Institute of Health grant R35 NS105038 (D.J.).

## Author contributions

Conceptualization, L.Y., N.S.C, and D.J.; Methodology: L.Y.; Investigation: L.Y.; Writing -Original Draft: L.Y. and D.J.; Writing - Review & Editing: L.Y., N.S.C, and D.J.; Visualization: L.Y.; Supervision: D.J.; Funding acquisition: L.Y. and D.J..

## Materials and Methods

### EXPERIMENTAL MODELS AND SUBJECT DETAILS

#### Mice

All experiments performed in this study were approved by the Institutional Animal Care and Use Committee (IACUC) of University of California San Francisco (UCSF). Experiments followed the ethical guidelines outlined in the NIH Guide for the care and use of laboratory animals. Tg(Trpv1-EGFP)MA208Gsat/Mmcd transgenic mouse line was generated by the MMRRC (#033029-UCD). Ndufs4fl/fl was acquired from The Jackson Laboratory.52 Uqcrq fl/fl was made as previously described.53 Trpv1Cre was made as previously described and deposited at The Jackson Laborary.54 Mice were raised under 12:12 light-dark cycles with ad libitum access to food and water. Both males and females were used in this study and no difference was noticed between them. No notable differences in body size were observed across genotypes.

#### Primary adult mouse DRG culture

DRG neurons were dissected from adult mice of both genders between 6 – 20 weeks of age. Dissected DRGs were collected into Leibovitz’s L-15 medium (Thermo Fischer Scientific) on ice and dissociated by two steps of enzymatic digestions. DRGs were digested in 40 U/ml papain (Worthington Biochemical) for 15 min at 37°C in a calcium- and magnesium-free (CMF) HBSS (UCSF Media Productions), followed by a second digest with 3 mg/ml collagenase (Worthington Biochemical), 4 mg/ml dispase II (Sigma-Aldrich), and 0.5 mg/ml DNaseI (Sigma-Aldrich) in CMF HBSS. Enzymes were deactivated by the addition of 10% horse serum (UCSF media Productions) and triturated in 0.5 mg/ml DNaseI in CMF HBSS. DRG neurons were enriched from debris and support cells by passing through a 100 µM strainer and additional purification through a Percoll gradient was performed for long-term cultures. Specially, dissociated cells were centrifuged through a discontinued Percoll (Sigma-Aldrich) gradient consisting of 1 ml 12.5% Percoll in 0.5 mg/ml DNaseI in CMF HBSS, 2 ml 25% Percoll in 0.5 mg/ml DNaseI in CMF HBSS, and 1 ml 50% Percoll in CMF HBSS for 20 min at 800 xg with slow deceleration. The interphase between 25% and 50% Percoll was collected and washed with 10 ml CMF HBSS twice by centrifuging at 300xg for 5 min. All centrifugation steps were performed at 4 °C. DRG neurons were cultured in Ham’s F-12 Nutrient Mix which contains 1 mM pyruvate in its formulation (Thermo Fischer Scientific) with 10% Horse Serum, 50 ng/ml Nerve Growth Factor 7S (Millipore Sigma), 5 µM FdU (Millipore Sigma), 5 µM uridine (Millipore Sigma) (DRG media), on Poly-D-lysine (Sigma-Aldrich) and laminin (Invitrogen) coated surfaces. For experiments that compare TRPV1+ and TRPV1-populations, 50 ng/ml BNDF (STEMCELL technologies) was added to the DRG media. DRG culture is fed every 2 days with fresh DRG media.

#### Generation and culture of immortalized cell lines

All cell culture were maintained at 37 °C with 5% CO2 in sterile humidified incubators. HEK293T cells were cultured in DMEM (UCSF Media Productions) with 10% Bovine Calf Serum (Hyclone). K562 cells were cultured in RPMI media with 10% Fetal Bovine Serum (FBS, Peak Serum) and 1% Pen strep (UCSF Media Production). Wildtype and MCU KO MEF cells were generous gifts from Dr. Yuriy Kirichok45 and were cultured in DMEM with 10% FBS. TRPV1 expressing stable cell lines of HEK293T, K562, and MEF were generated from parental cell lines transduced with lentivirus carrying TRPV1 with a fluorescent marker. TRPV1 expressing cells were enriched by Fluorescence Activated Sorting at UCSF Laboratory for Cell Analysis (LCA) using SONY SH800 (Sony Biotechnology) and single cell clones that express the fluorescent marker was sorted on 96 well plates. Sorted clones were further functionally selected based on their response to capsaicin by FURA imaging. K562 clones used for the CRISPRi screen was derived from dCas9-KRAS stable cells (parental)21 and the expression of dCas9-KRAS in TRPV1 expressing clones was also confirmed by transducing with sgRNAs targeting CD81 and confirmed with surface staining of APC anti-human CD81 antibody (bioLegend) followed by flow cytometry on an Attune NxT flow cytometer (Thermo Fisher Scientific) at UCSF LCA. Stable knockdown clones were obtained after lentiviral transduction of individual targeting and NT sgRNAs followed by puromycin (2 µg/ml, Thermo Fisher Scientific) selection. Expression of sgRNA containing vector was confirmed by flow cytometry of the fluorescent marker.

### METHOD DETAILS

#### Compound treatment

Small molecules inhibitors of cell death or mitochondria functions were reconstituted with DMSO with the following exceptions: CP-456733 and DFO were dissolved in water. Cells were treated as detailed below, all concentrations are of the final solution. Figure1A: Survival of TRPV1+ HEK293T cells after exposure to DMSO or 3 µM capsaicin for 5 hours in 0 or 2 mM EGTA (left) or in Ringer’s solutions without Ca2+ or Mg2+ or with NMDG+ substituting Na+ (right) was measured and normalized to vehicle controls. Figure1I: Survival of knockdown clones after 24-hour treatment of DMSO or 0.2 µM capsaicin was measured. Figure1K: Survival of TRPV1+ K562 cells that received pretreatment of PierA for 72 hours or AA for 48 hours prior to a 24-hour exposure to 0.4 µM capsaicin was measured and normalized to vehicle conditions (not shown). Figure 2: When indicated in (A-C and D-G), TRPV1+ K562 cells were pretreated with 4 nM PierA for 72h, or 10 nM AA for 48 hours, and supplemented with 1 mM pyruvate (Pyr) and 50 µg/ml uridine (Uri). Figure 2D-G: 24-hour treatment of capsaicin at D: 0.4 µM; E: 0.1 µM; F and G: 0.2 µM. Figure 3A-B: K562 cells were pretreated with either vehicle or 4 nM PierA supplemented with 1 mM pyruvate and 50 µg/ml uridine for 3 days, then treated with vehicle or 0.4 µM capsaicin for 4 hours. Figure 3Cii: 1 µM capsaicin for 24 hours. Figure 4B,C: 50 µM M40403 for 30 minutes followed by HyPer7 assay (B) or a 5-hour exposure of 6.25 µM capsaicin (C). Figure 4E: pretreating cells with 40 nM PierA or 20 nM AA supplemented with 1 mM pyruvate and 50 µg/ml uridine for 72 hours prior to HyPer7 assay. Figure 4F,G: pretreat cells with 15 µM S1QEL1.1, 50 µM S3QEL2, or vehicle for 6 hours followed by HyPer7 assay (F) or an overnight exposure of 0.4 µM capsaicin (G). Figure 5C,D: 10 µM ionomycin with 10 mM CaCl2 for 2 hours. Figure 5C: 3-day treatment of 400 nM PierA or 200 nM AA. Figure 5E: 10 µM mitoPQ for 48 hours or 3.3 µM for 72 hours. Figure 6C: 100 µM capsaicin for 24 hours. Figure 6F: 20 µM or 2 µM capsaicin for 24 hours.

#### Time-lapse imaging

For capsaicin-induced toxicity, cultured primary sensory neurons or transfected HEK293T cells were imaged overnight in a temperature, humidity, and CO2 controlled chamber on a Nikon inverted widefield fluorescent microscope at the Center for Advanced Light Microscopy at UCSF. An hour of baseline video was taken prior to the addition of capsaicin (Sigma-Aldrich) or vehicle treatments. Images were acquired every 10 min at multiple locations of the chambered slide. For ionomycin-induced toxicity, primary DRG culture was imaged at 5 min / frame at multiple locations in duplicated wells. After 15 minutes (3 frames), 10 µM ionomycin or DMSO with 10 mM CaCl2 were added and imaged for another 2 hours. The number of TRPV1-EGFP+ or TRPV1-EGFP-neurons survived treatments were counted manually, and statistics were tabulated using Excel (Microsoft) and graphed using Prism (GraphPad).

#### Molecular Cloning

Human TRPV1 cDNA was subcloned into pLenti IRES mApple vector (generous gift from Dr. Martin Kampmann at UCSF), which was used to generate TRPV1+ immortalized cell lines. Mouse Pgc1b (Addgene1031) and mouse Esrra (Addgene 172152) were subcloned into pLenti IRES mApple vector for overexpression experiments. Individual sgRNAs (top scoring targeting sgRNAs or NT controls) were cloned from custom oligos (Integrated DNA Technologies) using an annealing protocol (https://weissman.wi.mit.edu/crispr/) into pU6+ Ef1alpha Puro-T2A-GFP (Addgene 111596).

#### Lentiviral production, transductions, and storage

Early passage HEK293T cells were used to package lentivirus: they were transfected using Transit Lenti transfection reagent (Mirus Bio) and Opti-MEM (GIBCO), boosted with ViralBoost (Alstem, Inc) 24 hours after transfection, and harvested 48 hours post-transfection. K562 cells were transduced by “spinfection” for 2 hours (https://weissman.wi.mit.edu/crispria_cell_line_primer/). For transduction of neurons, harvested virus was concentrated 7-fold by precipitating with Lentivirus Precipitation Solution (Alstem, Inc) and reconstituting with DRG media. DRG culture was transduced on DIV2, fed every 2 days, and treated with capsaicin or vehicle on DIV9. Virus was stored at -80 °C and used within 2 months.

#### CRISPRi screen

A genome-wide single-guide RNA (sgRNA) library hCRISPRi-v2 top522 was packaged into lentivirus as previously described (https://weissman.wi.mit.edu/crispr/). TRPV1+ CRISPRi K562 cells were transduced with library virus at a multiplicity of infection (MOI) <1 (22% transduction efficiency 2 days post-transduction) and a coverage of > 1000 cells/sgRNA, which was maintained throughout the experiment. Replicates were maintained separately in spinner flasks for the course of the screen. Two days post-transduction, cells were selected with 0.75 µg/ml puromycin for 2 days to reach 80-90% transduced cells. After 24 hours of recovery from puromycin, a sample was collected from two biological replicates (T0), and remaining samples were further split into groups that received no treatment (NT) or pulsed capsaicin treatment (CAP). Cells were counted and split daily over the course of 14 days. CAP group received pulsed treatment of 0.2 µM (LD50) capsaicin until CAP cells had undergone 5-6 fewer doublings than UT cells92. Genomic DNA was isolated from T0, NT, and CAP samples using NucleoSpin Blood XL kit (Macherey-Nagel) and prepared for sequencing on an Illumina HiSeq4000 as previously described21. Sequencing reads were aligned and counted using a previously described MAGeCK based pipeline91.

#### Viability assays for immortalized cell lines

Cell viability of immortalized cell lines was performed on 96 well plates according to manufacturers’ protocols and measured using a Biotek H4 plate reader (Agilent). For all experiments, triplicated or quadruplicated samples of each condition were used. Triplicated samples of known number of cells counted by Countess II (Thermo Fisher Scientific) under each condition (parental, TRPV1+, or KD cell lines; vehicle or highest concentration of pretreatment drugs) were created on the same plate of each experiment. Estimated cell numbers were calculated when they are within linear range of the standards of the corresponding conditions. HEK293T cells were plated on PDL-coated 96 well plates with black walls, 5 hours after capsaicin treatment, cell numbers were estimated using alamarBlue (Thermo Fisher Scientific) fluorescence (Figure 1A, S2A, S3C, S4C and S4I) or cyQUANT fluorescent staining (Thermo Fisher Scientific) to identify nuclei of live cells (Figure 4C, 4G and S4J). K562 cells were plated on 96 well plates with white walls and treated with capsaicin overnight. Cell Titer Glo (Promega) was used to estimate total cell number (Figure 1I, 1K, 2D-G, 3Cii, S2B-D, S3A, S3D-H, S3J and S3K). Lactate dehydrogenase (LDH) assay (Thermo Fisher Scientific) was performed on the media supernatant taken after an overnight capsaicin treatment (Figure S3I). Data obtained from the plate reader was analyzed using Excel (Microsoft) and graphed with Prism 10 (GraphPad Software).

#### Immunoblotting

Cell lysate was extracted with PBST (0.5% Triton) with protease inhibitor cocktail (Roche) at 4°C for 30 min and supernatant was taken after centrifuging at 4°C 300g for 5 min. Proteins were resolved on 5-15% gels (bioRAD) and transferred onto PVDF membrane (0.2um). For small molecular proteins, blots were fixed in 4% PFA for 30 min at RT, prior to blocking. This is followed by standard immunoblotting procedures, and developed with ECL plus, ECL femto (Thermo Fisher Scientific), and imaged using a ChemiDoc Imager (Bio-Rad Laboratories).

#### RNAseq

TRPV1+ K562 cells were pretreated with vehicle or 4 nM PierA for 72 hours in the presence of 1 mM pyruvate and 50 µg/ml uridine followed by vehicle or 0.4 µM capsaicin treatment for 4 hours. There were 3 technical replicates of each condition, each replicate used 1.6 million cells. RNA was extracted using RNeasy Micro kit (Qiagen) and all replicates had RIN = 10 (Agilent Bioanalyzer). Library preparation, sequencing (Ilumina), and sequence alignment were done by Novogene Corporation Inc. Differential expression analyses, data visualization, and cross-referencing to CRISPRi screen was done using R. Reanalysis of published dataset (GEO: GSE131230)^49^ was done using R.

#### Calcium imaging and analysis

FURA-2AM imaging was performed as preciously described93. Briefly, cells were washed with Ringer’s solution (140 mM NaCl, 5 mM KCl, 2 mM MgCl2, 2 mM CaCl2, 10 mM HEPES and 10 mM glucose, pH 7.4 with NaOH; 290–300 mOsm kg−1) once and loaded with 10 μg/ml Fura2-AM (Thermo Fisher Scientific) in Ringer’s solution containing 0.02% Pluronic F-127 (Thermo Fisher Scientific) at room temperature in the dark for 1 hour. Following another wash with Ringer’s solution, they were immediately imaged on an inverted microscope with 340 and 380 nm excitation (Sutter, Lambda LS illuminator) at a time interval of 2s. MetaFluor (Molecular Devices) was used for image acquisition and analyses. For FURA-2AM loaded K562 cells, 50 random cells were selected from each video to analyze their response (emission ratio of 340/380) to 33 nM capsaicin and 10 µM ionomycin. Baseline (F0) was calculated as the average of 340/380 ratio prior to drug application, and ΔF was calculated as the stable 340/380 emission ratio after drug applications subtracted by the F0. Response to capsaicin was presented as a fraction of maximum response to ionomycin: ΔFCap / ΔFIonomycin. For FURA-2AM loaded DRG culture used to determine TRPV1+%, Ringer’s was added to control for activation of mechanosensitive neurons, followed by the addition of 10 µM capsaicin, and finally high-K+ Ringer’s solution (same as Ringer’s except 5 mM NaCl and 140 mM KCl) to identify neurons.

#### HyPer7 imaging and analysis

Stable TRPV1 expressing HEK293T cells were transfected with mitochondrial-targeted HyPer7 (ref, Addgene). Live cells were imaged on an inverted Nikon Ti microscope at 37°C with CO2. HyPer7 was excited at 380 nm and 480 nm, and emission was collected at 525 nm over 45 min with 1 min intervals using a 40x objective. In a typical independent experiment, 5 videos are obtained from each well, and each experimental conditions were duplicated in two wells. Videos were analyzed using the NIS-Elements (v5.4) to obtain HyPer7 ratio (F = fluorescent intensity of emission signals excited by 480nm/380nm) of each cell over time. Response to 20 µM capsaicin and 200 µM H_2_O_2_ was calculated by R (v2023.03.0+386) using the formula ΔF/F0 where ΔF is the difference between HyPer7 ratio at each time points and the average of the baseline timepoints (before capsaicin addition) and plotted using Prism 10. Summary data presented were from at least 3 independent experiments.

#### MitoSOX assay

Parental or TRPV1+ HEK293T cells were plated on 12-well plates and grew to 70% confluent on the day of capsaicin treatment. A fresh aliquot of lyophilized mitoSOX (Thermo Scientific) was reconstituted in DMSO and used at the final concentration of 50 µM during the last 20 min of capsaicin or vehicle treatment. Cells were harvested on ice, centrifuged at 300xg for 5 min, and cell pellets were washed with PBS. Wash cell pellets were resuspended in PBS with 10% FBS and analyzed on an Attune NxT flow cytometer (Thermo Fisher Scientific). Flow cytometry data was analyzed using FlowJo (FlowJo LLC) and Prism (GraphPad Software). Gating was determined using no mitoSOX and vehicle treatment controls and used for all samples of the same experiment.

#### Neurotoxicity assays

DRG culture was plated on laminin (Invitrogen) coated 18-well chamber slides (ibidi GmbH, Cat No. 81816). On or after DIV2, primary DRG culture media was replaced with fresh media containing vehicle or compound treatments of indicated concentrations and durations. Capsaicin or vehicle treatments were supplemented with 2 mM CaCl2 and incubated overnight. After which, cells were washed with PBS, replaced with DRG media, and recovered for at least 24 hours before performing FURA imaging to determine the percentage of TRPV1+ neurons (TRPV1+%) in each condition. TRPV1+% was calculated from the number of neurons responded to 10 µM capsaicin out of the number of neurons responded to high extracellular K+. Percentage of TRPV1+ neuron that survived capsaicin treatment (X) was calculated from the TRPV1+% after vehicle (Y) or capsaicin treatment (Z) under the same genetic or pretreatment conditions, where X = (Z(100-Y))⁄(Y(100-Z))% (Figure S6). Videos were collected from at least 3 technical replicates (n) of each condition per independent experiment (N). Technical replicates from the same condition of each independent experiment were aggregated into a single data point. This assay was devised because plate reader assays were unsuitable due to the small quantity of DRG neurons per animal, heterogeneity of different kinds of neurons, and the presence of proliferating support cells. We also found assays, such as propidium iodide, that stain nuclei of dead cells inaccurate, as nuclei are often missing from ruptured plasma membrane of capsaicin-evoked necrotic cells. Due to variable percentage of TRPV1+ and baseline toxicity across biological replicates, we paired our analyses within each independent experiment.

#### Immunohistochemistry

TGs and DRGs were acutely harvested into ice-cold L-15 medium and transferred into 4% paraformaldehyde for overnight fixation. After washes with phosphate-buffered saline (PBS, Quality Biological, 119-069-491), the ganglia were allowed to settle in 30% sucrose (SigmaAldrich, S7903) at 4°C and were then cryopreserved in Tissue-Tek O.C.T. Compound (Sakura Finetek USA) for sectioning at 10 μm thickness. Sections were washed with PBST (i.e., PBS containing 0.3% Triton X-100 [Sigma-Aldrich, T8787]) and incubated with blocking buffer (PBST containing 10% bovine serum albumin (BSA) (Sigma-Aldrich A2153) and 10% normal goat serum (NGS) (Thermo Fisher Scientific, 16-210-064) at room temperature for 1 hr. Subsequently, sections were incubated with primary antibodies in PBST 1% BSA 1% NGS at 4°C overnight. Following 3 washes, the sections were incubated with secondary antibodies (Thermo Fisher Scientific) in blocking buffer at room temperature for 1 hr. Finally, the sections were washed and mounted with Fluoromount-G (SouthernBiotech, 0100-01). Images were taken on a Nikon inverted widefield fluorescent microscope (UCSF Center for Advanced Light Microscopy). Same parameters were used to collect images of the same experiment. To quantify ETC expression, mean fluorescent intensities of TRPV1+ cells and their closest neighboring TRPV1-cells were measured using Fiji (54). Primary antibodies: rabbit anti-TRPV1 (Alomone Labs, ACC-030, 1:400), chicken anti-GFP (abcam, AB13970, 1:1000), rabbit anti-Ndufs4 (Millipore Sigma, HPA003884, 1:100), rabbit anti-Cox6 (Millipore Sigma HPA014295; 1:100).

---

**Figure S1.**
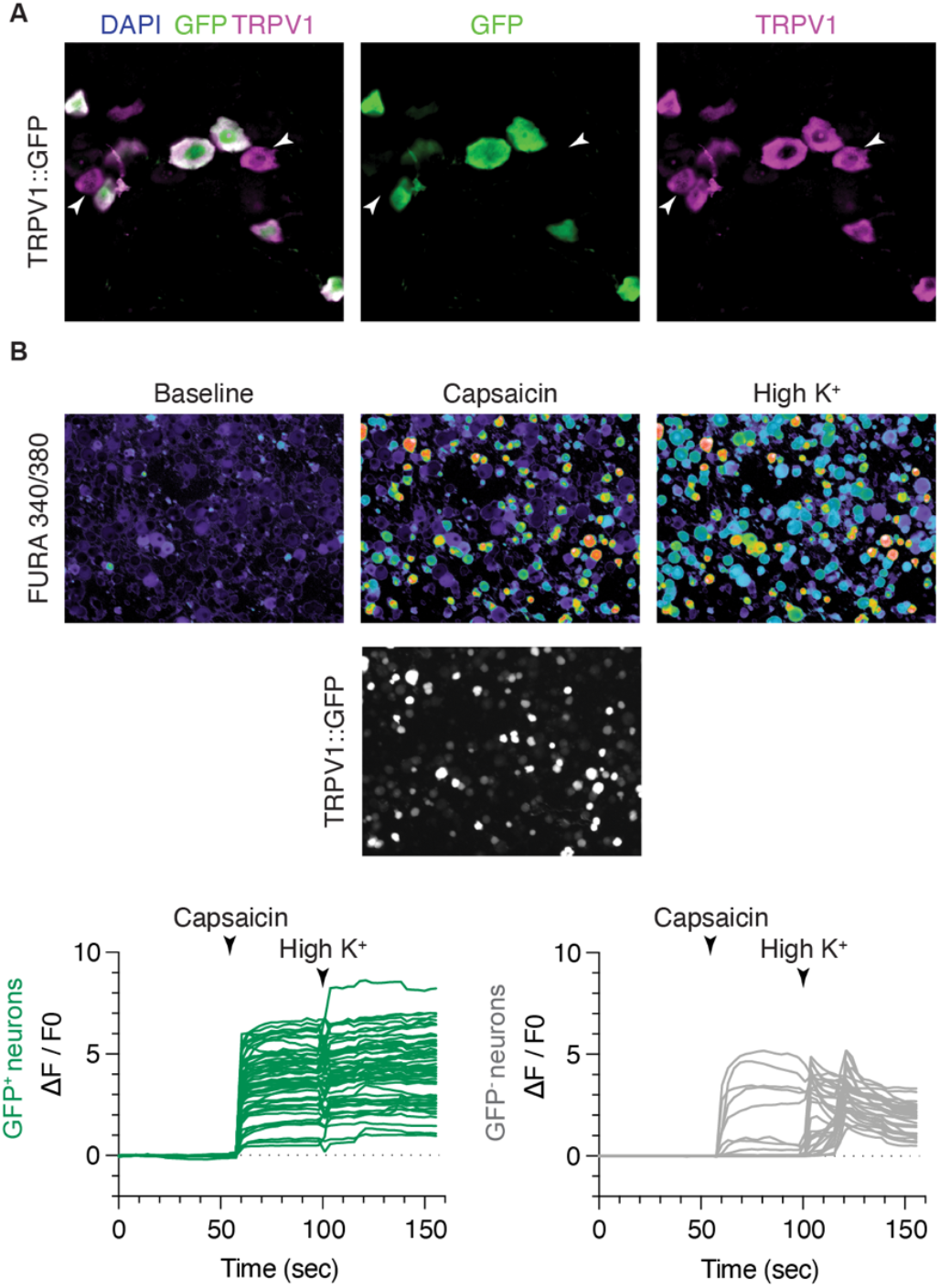
Characterization of the BAC transgenic reporter mouse line *Tg (Trpv1-EGFP) MA208Gsat / Mmcd*. (Related to Figure 1 and 5) Because the EGFP reporter is driven by a *Trpv1* promotor on a Bacterial Artificial Chromosome (BAC), we characterized GFP^+^ versus GFP^-^ neurons for their expression of TRPV1. **(A)** Immunostaining of trigeminal ganglia (TG) sections with TRPV1 and GFP antibodies. All GFP^+^ cells were also positive for TRPV1 staining. However, a small fraction of TRPV1^+^ cells were not positive for GFP staining (arrows). **(B)** Dissociated DRG culture was loaded with FURA-2AM dye. Calcium influx was recorded as an increase of 340/380 ratio. Consistent with (A), all GFP^+^ cells showed calcium response to capsaicin, and a small fraction of GFP^-^ cells was also responsive to capsaicin. High extracellular potassium (high K^+^) was used to reveal all neurons. In summary, both histological and physiological evaluations of the *Tg(Trpv1-EGFP)MA208Gsat/Mmcd* mouse showed no false positives, although some false negatives were observed using both methods. For simplicity, GFP^+^ neurons from this line are referred to as TRPV1^+^ in Figure 1B, 5B, and 5D.

**Figure S2.**
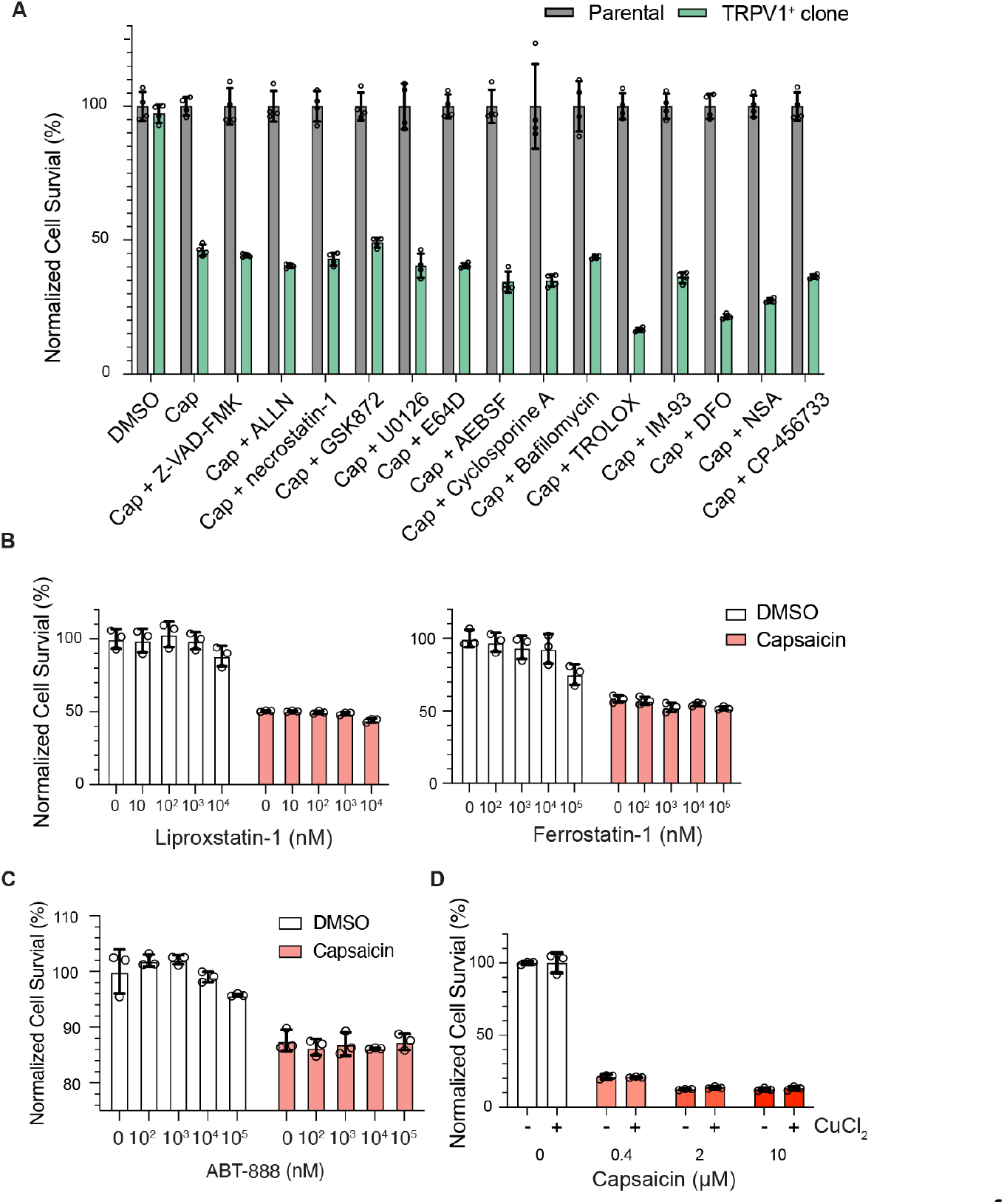
Characterization of capsaicin-induced cell death with small molecule inhibitors of cell death modalities. (Related to Figure 1) **(A)** Parental or TRPV1^+^ HEK293T cells were treated with DMSO, 6 µM Capsaicin, or 6 µM Capsaicin with 45 µM Z-VAD-FMK (pan caspase and apoptosis inhibitor), 40 µM ALLN (calpain inhibitor), 20 µM necrostatin-1 (RIPK and RIPK dependent apoptosis inhibitor), 3.3 µM GSK872 (RIPK3 and necroptosis inhibitor), 52 µM U1026 (MAPK inhibitor), 80 µM E64D (cysteine protease and lysosome-dependent cell death inhibitor), 417 µM AEBSF (serine protease inhibitor), 33 µM cyclosporine A (calcineurin and mitochondrial permeability transition driven cell death inhibitor), 4 µM Bafilomycin (lysosomal V-ATPase and autophagy inhibitor), 6.25 mM 3-MA (autophagy inhibitor), 240 µM TROLOX (antioxidant and ferroptosis inhibitor), 25 µM IM-93 (ferroptosis and NETosis inhibitor), 100 µM DFO (deferoxamine: iron chelator and ferroptosis inhibitor), 173 µM NSA (necrosulfonamide inhibits GSDMD and pyroptosis as well as MLKL and necroptosis), and 188 µM CP-456733 (or MCC950 inhibits NLRP3 inflammasome and pyroptosis).^20^ Cell number was measured using AlamarBlue and normalized to DMSO treated parental line. Plate replicates n=4, representative of at least 3 independent experiments. **(B**,**C**,**D)** Ferroptosis, parthanatos, and copper induced cell death were examined using potent lipid peroxidation inhibitors Liproxstatin-1 and Ferrostatin-1 **(B)**,^87^ PARP-1 inhibitor ABT-888 **(C)**,^88^ and 10 µM extracellular CuCl2 to assess contribution of copper influx through TRPV1 **(D)**.^89^ TRPV1^+^ K562 cells pretreated with vehicle or serial dilutions of inhibitors for 30 min followed by 0.4 µM capsaicin or vehicle treatment for 24 hours. Cell number was measured using Cell Titer Glo and normalized to vehicle pretreated and vehicle treated samples. Plate replicates n=3. Representative of at least 2 independent experiments. Data are mean SD.

**Figure S3.**
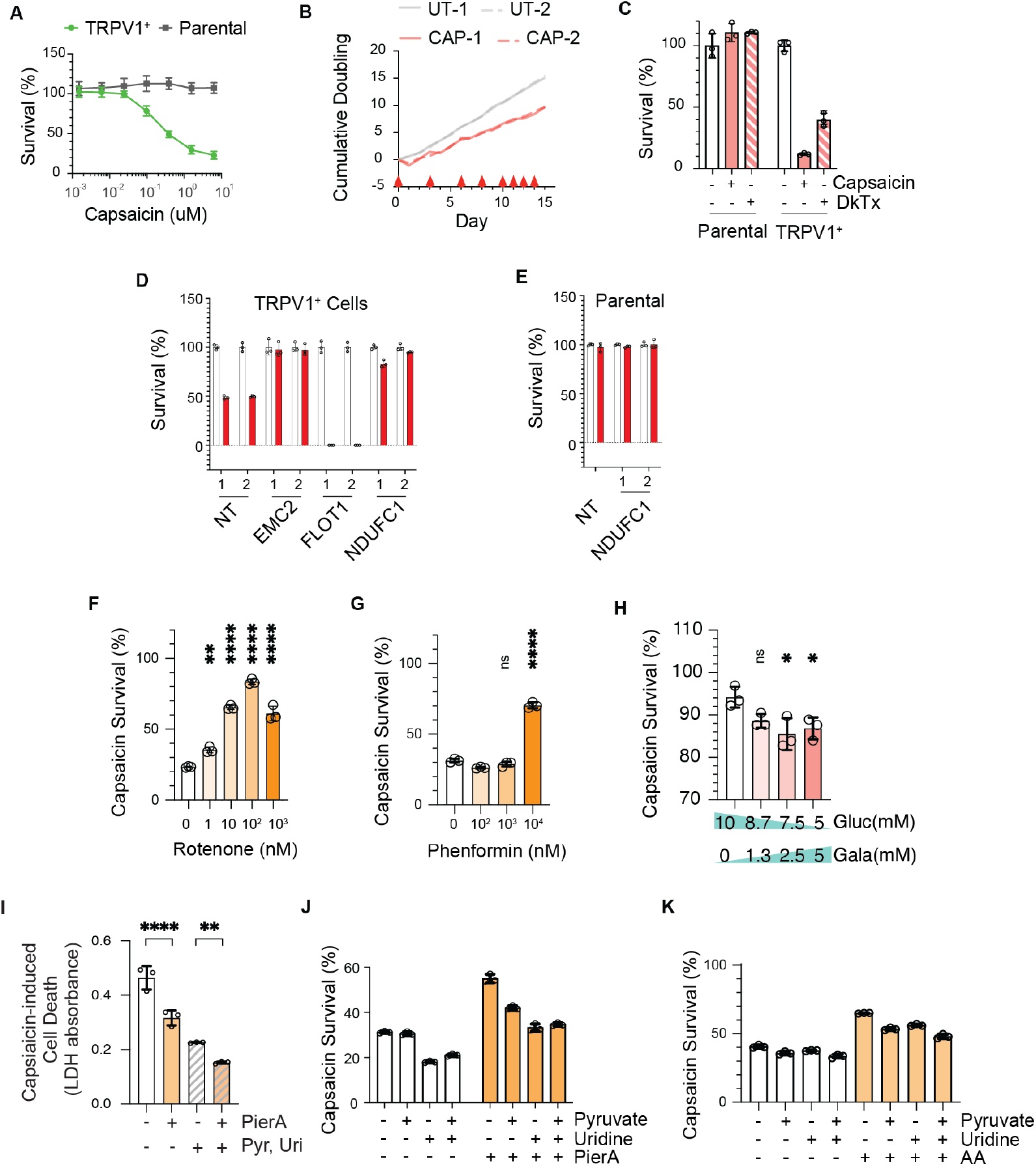
Additional information of the CRISPRi screen and characterization of the metabolic regulation of capsaicin-induced cell death. (Related to Figure 1,2) **(A)** Capsaicin LD50 characterization of TRPV1^+^ K562 dCas9-KRAB cells and the parental K562 dCas9-KRAB cells were determined using Cell Titer Glo. The capsaicin toxicity is specific to cells that express TRPV1. **(B)** Cumulative doubling of untreated (UT) and capsaicin (CAP) treated duplicates were calculated from daily cell density counted using Countess 2 Automated Cell Counter from day 0 to 14 when 5-6 doubling differences was achieved. Red arrow heads indicate pulse treatment of capsaicin (half lethal dose 50). **(C)** Survival of HEK293T cells (left) or TRPV1^+^ HEK293T cells (right) after 5-hour treatment of DMSO control, 3 µM capsaicin, or 3 µM DkTx was measured using alamarBlue and normalized to DMSO control. **(D**,**E)** Viability of TRPV1^+^ CRISPRi K562 cells (D) or CRISPRi K562 cells (E) that stably expressed each sgRNA were determined using Cell Titer Glo after 24-hour treatment of DMSO control or 0.2 µM capsaicin and normalized to DMSO controls of each stable KD line. Two sgRNAs of each target gene and non-targeting (NT) controls were used. (D) contains vehicle controls that is not shown in Fig. 1F. As capsaicin survival is always normalized to vehicle control of the same condition, vehicle controls are not shown in all other viability assays. **(F**,**G)** TRPV1^+^ K562 cells was pretreated with (F) rotenone or (G) phenformin for 72 hours and then treated with 0.4 µM capsaicin or vehicle control for 24 hours. Apoptosis was observed in samples pretreated with 0.1 and 1 µM rotenone prior to capsaicin exposure. **(H)** TRPV1^+^ K562 cells grew in RPMI media with indicated concentrations of glucose and galactose for 72 hours days and treated with 0.2 µM capsaicin or DMSO control. Cell number was measured using Cell Titer Glo and normalized to DMSO control (not shown) of each condition. **(I)** Necrosis was estimated by the amount of lactate dehydrogenase (LDH) in culture medium as an alternative to quantifying survivors as in (Figure 2E-2G), conditions were the same as (Figure 2F). **(J**,**K)** TRPV1^+^ K562 cells were pretreated with 4 nM PierA for 3 days (I), 10 nM AA for 2 days (J) with the indicated supplements (1 mM pyruvate or 50 µg/ml uridine) or vehicle controls. (See Figure 2F and 2G). Data are mean SD. N=3 of technical replicates representative of at least 3 independent experiments. (F-I) One-way ANOVA with Dunnett’s multiple comparisons test using veh (F,G,I) or no Galactose (H) as controls. * P < 0.05; ** P<0.01; **** P<0.0001.

**Figure S4.**
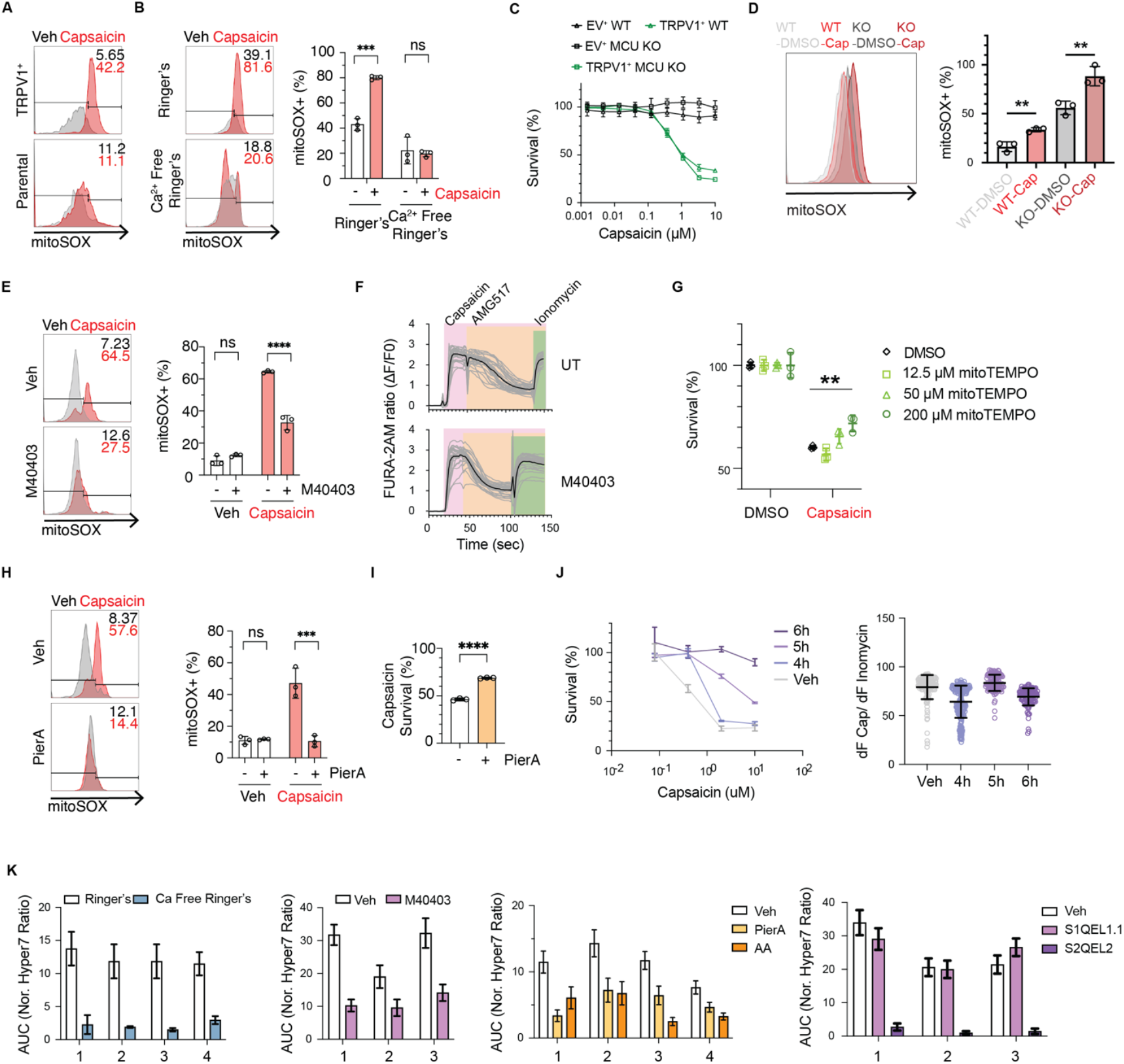
Additional characterization of capsaicin-evoked oxidative stress. (Related to Figure 4) **(A)** Mitochondrial superoxide (O2-) level was measured by flow cytometry after mitoSOX staining. For TRPV1^+^ HEK293T, percentage of mitoSOX^+^ cells increased after 30 min 10 µM capsaicin treatment comparing to vehicle controls, whereas little difference was observed for parental HEK293T cells. **(B)** TRPV1^+^ HEK293T cells treated with 10 µM capsaicin or vehicle in regular or calcium free Ringer’s solution for 1 hour. Left: representative mitoSOX histograms with gating frequencies; Right: percentage of mitoSOX^+^ population from 3 biological replicates. **(C)** Lethal dose of capsaicin was determined for MEF cell clones that express empty vector (EV) or TRPV1 derived from wildtype (WT) or MCU knockout (KO) clones. Lack of MCU did not rescue capsaicin-induced cell death. **(D)** TRPV1-expressing MEF clones of either WT or MCU KO background were treated with 30 min of 10 µM capsaicin or vehicle. Increases of mitoSOX fluorescence in response to capsaicin treatment was observed of both clones. Although there is a difference between WT and KO clones under the DMSO conditions, we cannot conclude such difference is due to KO as they were single cell clones derived from different parental lines. Representative histograms on the left; summary of triplicates on the right. **(E)** TRPV1^+^ HEK293T cells pretreated with 50 µM M40403 or vehicle for 30 min, followed by 30 min treatment of 10 µM capsaicin or vehicle and mitochondrial O2-level was measured by mitoSOX assay. **(F)** Pretreatment of M40403 does not change capsaicin evoked calcium influx. Dynamics of calcium influx was recorded of HEK293T TRPV1 clones that were (i) untreated or (ii) pretreated with 50 µM M40403 for 30 min. Calcium response to 0.1 µM capsaicin, 2 µM AMG517 (TRPV1 antagonist), and 1 µM ionomycin were recorded as 340/380 ratio of FURA-2AM fluorescence. Randomly selected 30 cells (grey) and the average (black) were shown above. No significant difference in capsaicin-induced calcium influx was observed between untreated and M40403 treated cells. Experiments showed here are representative of 3 independent repeats. **(G)** TRPV1^+^ K562 cells were pretreated with indicated concentrations of mitoTEMPO for 30 min prior to a 24hour incubation with 1 µM capsaicin or vehicle. **(H**,**I)** TRPV1^+^ HEK293T cells pretreated with 20 nM PierA or vehicle and supplemented with 2 mM pyruvate and 0.1 mg/ml uridine for 72 hours. Subsequently, (H) mitochondrial O2-level was measured after 30 min treatment of 10 µM capsaicin or vehicle; (I) viability was measured after 6-hour exposure to 100 µM Capsaicin or vehicle. **(J)** TRPV1^+^ HEK293T cells pretreated with S3QEL2 for 4,5,6 hours or vehicle for 6 hours, followed by viability assay (left, using cyQUANT) or calcium response assay (right, using FURA2-AM). **(K)** AUC quantifications of each independent experimental replicate summarized in Figure 4. Data are mean SD. **(B**,**E**,**H)**: One-way ANOVA Tukey’s multiple comparisons test. **(D**,**I)**: 2-tailed unpaired t-test. **(G)** One-way ANOVA with Tukey’s multiple comparisons test. 0 vs 200 µM mitoTEMPO. ** P<0.01, *** P<0.001, **** P<0.0001.

**Figure S5.**
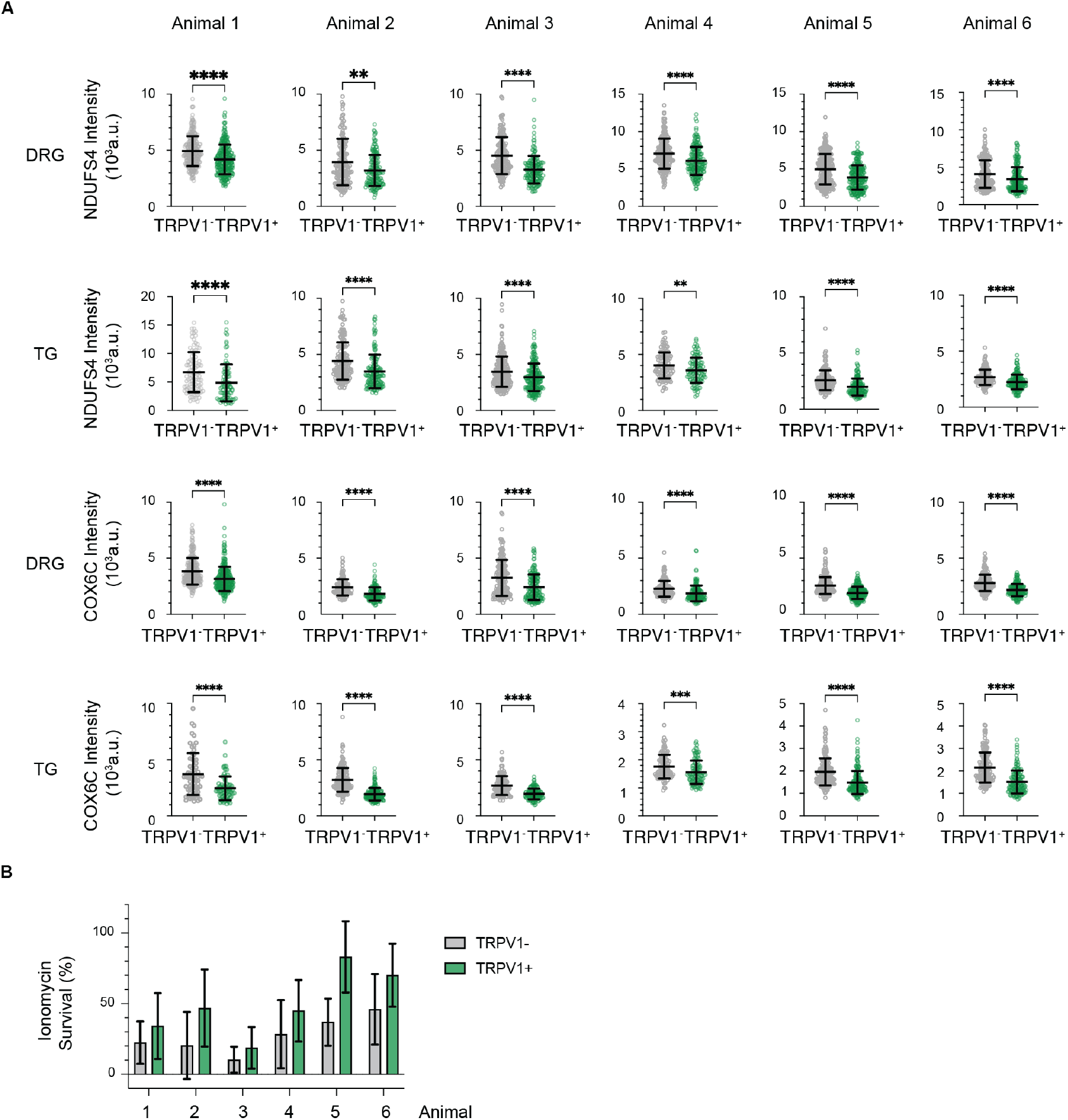
Expanded data related to Figure 5. (Related to Figure 5) **(A)** Quantification of the staining intensities of NDUFS4 or COX6C in TRPV1^+^ versus TRPV1^-^ in DRG or TG sections from each of the 6 animals. Each dot represents a cell from at least 5 histological sections of indicated tissue per animal. **(B)** Relative sensitivity to ionomycin of TRPV1(GFP)^+^ versus TRPV1(GFP)^-^ neurons quantified from 6 animals and at least 10 videos per animal.

**Figure S6.**
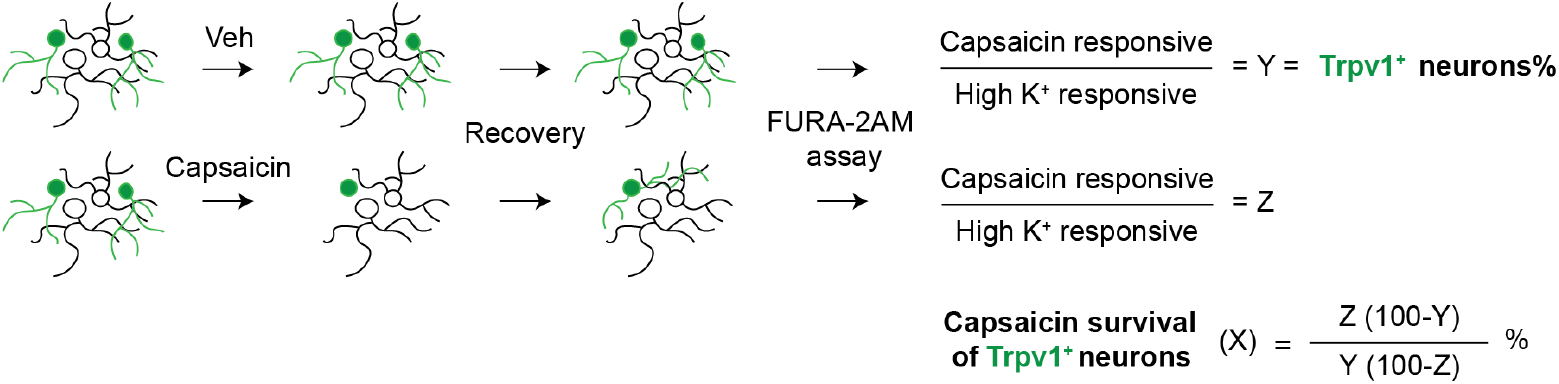
Schematic representation of capsaicin survival assay of DRG culture. (Related to Figure 6) More details in Methods. Briefly, the percentage of TRPV1^+^ neurons that survived capsaicin treatment (X) was calculated from the percentage of capsaicin responding neurons in vehicle (Y) or capsaicin (Z) treated samples.

